# Homogeneous multifocal excitation for high-throughput super-resolution imaging

**DOI:** 10.1101/2020.01.08.895565

**Authors:** Dora Mahecic, Davide Gambarotto, Kyle M. Douglass, Denis Fortun, Niccoló Banterle, Maeva Le Guennec, Khalid Ibrahim, Pierre Gönczy, Virginie Hamel, Paul Guichard, Suliana Manley

## Abstract

Super-resolution microscopies, which allow features below the diffraction limit to be resolved, have become an established tool in biological research. However, imaging throughput remains a major bottleneck in using them for quantitative biology, which requires large datasets to overcome the noise of the imaging itself and to capture the variability inherent to biological processes. Here, we develop a multi-focal flat illumination for field independent imaging (mfFIFI) module, and integrate it into an instant structured illumination microscope (iSIM). Our instrument extends the field of view (FOV) to >100×100 µm^2^ without compromising image quality, and maintains high-speed (100 Hz), multi-color, volumetric imaging at double the diffraction-limited resolution. We further extend the effective FOV by stitching multiple adjacent images together to perform fast live-cell super-resolution imaging of dozens of cells. Finally, we combine our flat-fielded iSIM setup with ultrastructure expansion microscopy (U-ExM) to collect 3D images of hundreds of centrioles in human cells, as well as of thousands of purified *Chlamydomonas reinhardtii* centrioles per hour at an effective resolution of ∼35 nm. We apply classification and particle averaging to these large datasets, allowing us to map the 3D organization of post-translational modifications of centriolar microtubules, revealing differences in their coverage and positioning.

## Introduction

Super-resolution fluorescence techniques have enabled optical imaging beyond the diffraction limit. Two main illumination strategies are used to achieve super-resolution: wide-field illumination is typically used for single-molecule localization microscopies (SMLM) such as STORM^1^ and PALM^2^, while patterned illumination is typically used for methods such as structured illumination microscopy (SIM)^3,4^ and stimulated emission depletion (STED)^5^. Initial implementations of these methods were relatively slow, in part due to the use of the time domain to separate single molecules (SMLM), the need to collect multiple images to cover Fourier space (SIM), or the scanning of a reduced excitation volume (STED). In both wide-field and patterned illumination, the imaging throughput – defined as the area imaged per time – can be increased by parallelizing the acquisition, as long as the necessary illumination can be maintained over a larger surface in the sample plane. Extending the array of patterned excitation has been used to increase the speed or FOV of confocal^6^, STED^7,8^ and SIM imaging^9^, but ensuring the quality of the extended pattern across a larger FOV remains a limiting factor. For SMLM, flat-fielding approaches made it possible to extend super-resolution imaging over ∼100×100 µm^2^ FOVs^10,11^, but their transfer to patterned illumination remains limited.

An entirely different approach to effectively achieve super-resolution modifies sample preparation rather than image acquisition. By physically increasing the size of the specimen in an isotropic manner via expansion microscopy^12^ (ExM), overall imaging throughput can be increased by opting for faster diffraction-limited microscopes^12–14^. Alternatively, combining ExM with existing super-resolution microscopies allows even further improvement in resolution^14–18^, since the resolvable scale is reduced by the expansion factor. However, expansion exacerbates limitations in imaging throughput since the effective FOV size is divided by the expansion factor along each dimension^19^. Therefore, fast, parallelized, super-resolution techniques with large FOVs present an advantage for imaging ExM samples, among other applications.

The instant structured illumination microscope (iSIM)^9^ uses parallelization in the form of multifocal excitation and optical image processing to achieve a two-fold improvement in resolution, at frame rates reaching 100 Hz. While this gives the iSIM good potential for high-throughput super-resolution imaging, spatially varying illumination produces a spatially dependent signal to noise ratio while restricting the imaging FOV to ∼50×50 µm^2^. Although solutions for producing homogeneous multi-focal excitation exist using diffractive optical elements^20^, spatial light modulators (SLM)^21^, multi-mode fibers^22^ or beam splitters^23^, each suffers from some combination of a limited number of excitation spots^20,21,23^, heterogeneity^21,23^ or low transmission efficiency^20^.

In this work, we develop a flat-fielding element for multi-focal excitation, dubbed mfFIFI. Our design is based on the Koehler integrator^24–26^ and ensures that the homogeneity, pitch and size of the excitation spots are optimized. We integrate mfFIFI into an iSIM to engineer an instrument capable of 3D multi-color imaging with uniform quality and doubled resolution over a 100×100 µm^2^ FOV. We further parallelize the acquisition across multiple FOVs, stitching them together to create seamless montages of tens of super-resolved living cells within seconds, enabled by the homogeneous illumination. Finally, we combine the mfFIFI iSIM with ultrastructure expansion microscopy (U-ExM)^31^ to perform high-throughput super-resolution imaging with an effective resolution of ∼35 nm. We use this to collect 3D images of hundreds of expanded centrioles in human cells, and thousands of expanded purified *Chlamydomonas reinhardtii* centrioles. These particles are averaged to reconstruct maps of post-translational modifications (PTMs) of centriolar tubulin, revealing differences in the spatial organization of acetylation, monoglutamylation and polyglutamylation.

## Results

### Design and implementation of homogeneous multifocal excitation

Generating a uniform irradiance over a large FOV while retaining the highest achievable resolution requires the following properties of the excitation focal spots in a multi-focal microscope: 1) homogeneous intensity, 2) constant pitch and 3) diffraction-limited size across the field of view. Existing methods for generating uniform excitation are often limited to producing a low number of excitation spots, with residual heterogeneity and low transmission efficiency. Thus, we set out to extend the Koehler integrator^24–26^, which averages over the spatial and angular distributions of the light source to homogenize the beam, for use with multifocal excitation.

Previously, multifocal excitation for the iSIM was generated by illuminating a microlens array (MLA) with a collimated Gaussian beam^6,9^. As expected, the non-homogeneous beam profile results in excitation points whose intensity depends on their location, a result which is recapitulated in a wave optics simulation (**Figure 1a**) (**Supplemental information**). Alternatively, illuminating the excitation MLA with a flat-top beam – such as that produced by a Koehler integrator – results in an array of multifocal excitation spots of homogeneous intensity (**Figure 1b**). This can be achieved by placing the excitation MLA at the front focal plane of the Fourier lens of the Koehler integrator, where the flat-top is focused (**Supplemental Figure 1a**). However, if the incident wavefront is not flat, the non-telecentric illumination of the excitation MLA produced by this configuration alters the pitch of the excitation spots in the focal plane of the excitation MLA (**Figure 1c**). This periodicity – usually determined by the pitch of the excitation MLA – must match the periodicity of additional components that are typically placed in conjugate image planes of confocal microscopes, such as a pinhole array. In the case of a mismatch of array elements, the light will be gradually occluded and decrease in intensity away from the center of the optical path. To solve this problem and achieve telecentric illumination of the excitation MLA, we place the Fourier lens of the Koehler integrator one focal length away from the second flat-fielding MLA^27^, resulting in a flat wavefront and thereby ensuring that the pitch of the excitation is conserved (**Figure 1d, Supplemental Figure 1b**). Additionally, adjustment of the distance between the second flat-fielding MLA and the Fourier lens can be used to finely adjust the resulting periodicity of the excitation spots to match that of the other optical elements (**Figure 1e, Supplemental Figure 2a-d**). This feature, not shared by refractive beam-shapers, allows us to optimize transmission efficiency.

**Figure 1.**
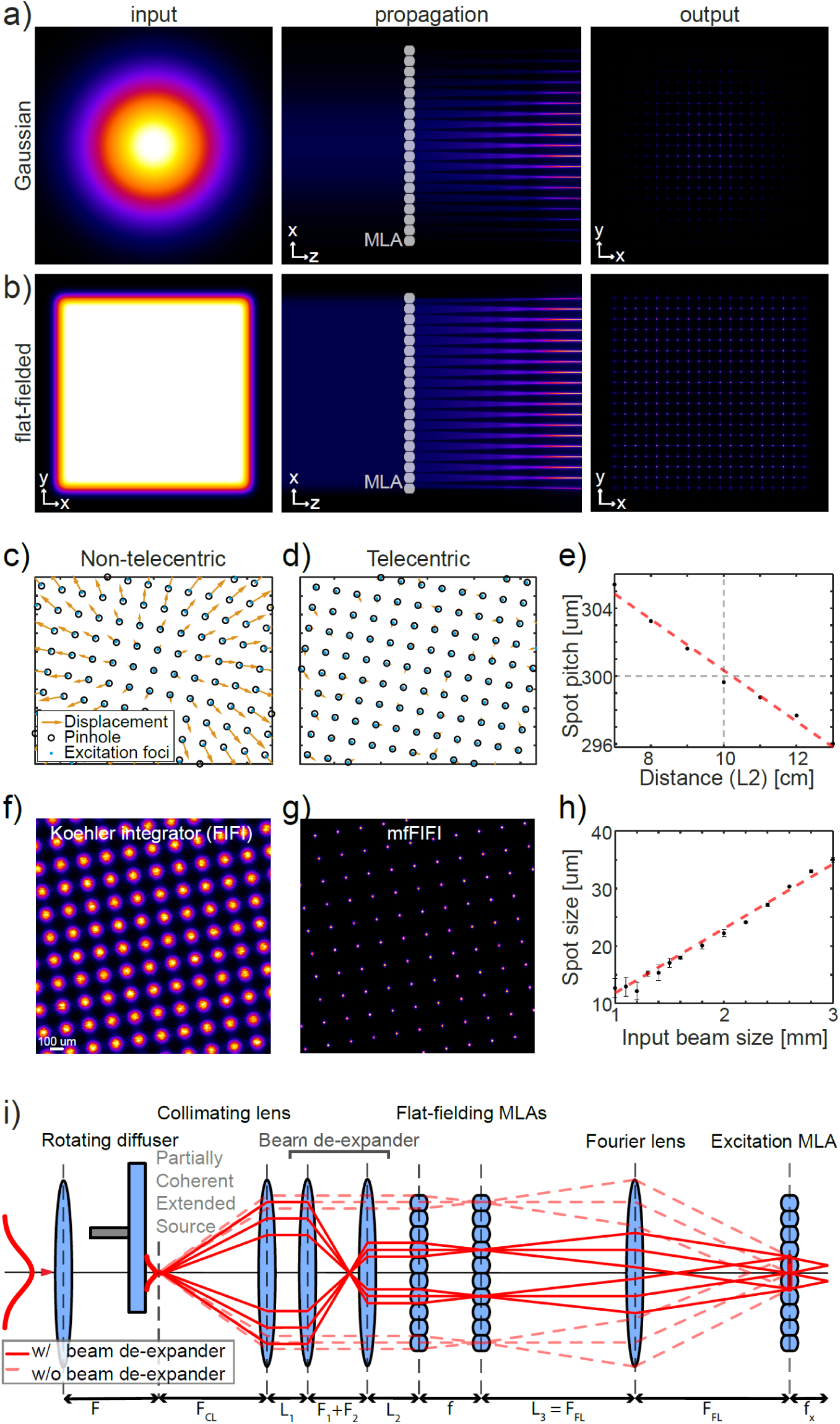
Design and features of the mfFIFI module. a,b) Wave optics simulation with (a) a Gaussian beam or (b) a flat-top beam used as input onto the excitation MLA, showing their propagation and their effect on the output intensity of the generated excitation spots. c,d) Experimental positions of the excitation spots (blue dots) overlaid on a conjugate pinhole array (black circles) and the displacement between the two (yellow arrows) for (c) non-telecentric and (d) telecentric illumination of the excitation MLA reveals a change in the pitch of excitation spots caused by non-telecentric illumination. Data are from separately acquired images of pinholes and excitation spots. e) Plot showing the dependence of the pitch of the excitation spots on the offset of the Fourier lens back focal plane to the second flat-fielding MLA. Dashed line represents zero offset on the x-axis and the ground truth set by the pitch of the MLA on the y-axis. f) Image of excitation spots generated with a Gaussian beam as input into the excitation MLA. Directly implementing the Koehler integrator as the input results in large excitation spots. g) Limiting the number of flat-fielding MLA channels used for beam homogenization partially recovers the size of the excitation spots as achieved using Gaussian excitation (a), while maintaining uniform intensity of the Koehler integrator (f). h) The spot size increases linearly with the diameter of the beam incident on the flat-fielding MLAs over the given range. i) Complete design and implementation of the extended Koehler integrator for multi-focal flat illumination for field independent imaging (mfFIFI).

Finally, in confocal microscopy and its variants such as iSIM, maximizing the achievable resolution requires focusing the excitation light to a diffraction-limited spot on the sample. This imposes a limitation on the spot size that should be generated in the excitation path. However, implementing a traditional Koehler integrator in the excitation path would produce large excitation spots due to the nature of the partially coherent extended source created by the rotating diffuser (**Figure 1e-f, Supplemental Figure 1**). In that case, the accessible improvement in resolution would not be based on diffraction-limited performance. A possible solution would be to introduce a pinhole array to mask the excitation spots that are focused onto the sample, but at a significant cost to the transmission efficiency.

In an alternative design, we find that – contrary to the typical Koehler integrator where illuminating a maximal number of flat-fielding microlenses is preferred^24–26^ – illuminating fewer microlenses offers a solution to control the excitation spot size and ensure diffraction-limited excitation at the sample (**Figure 1g,h**). Incorporating a beam expander to contract the light incident on the flat-fielding MLAs (referred to as a beam contractor henceforth, **Figure 1i, Supplemental Figure 1c**) allows us to tune the apparent size and angular distribution of the extended source. This shrinks the size of the excitation spots, while maintaining efficient light transmission and homogeneity across the excitation illumination. This configuration, which we call multifocal flat illumination for field-independent imaging (mfFIFI), meets all the above requirements for homogeneous multifocal excitation and is shown in **Figure 1i**.

These requirements can be described by geometrical optics and are a direct outcome of ray transfer matrix calculations (**Supplemental information, Supplemental Figure 2**), but the large parameter space makes the design and optimization of the mfFIFI module non-trivial. For example, beam contraction presents a trade-off between homogeneity of the multifocal excitation and the ability to achieve diffraction-limited excitation and requires careful optimization (**Supplemental information, Supplemental Figure 3h**). To facilitate the optimization and choice of components for efficient mfFIFI, we provide the main design equations and an extended version of our existing wave optics simulation platform^10^ (**Supplemental information, Supplemental Figure 3, Supplemental Table 1**).

### iSIM integration and performance characterization

We tested the performance of the mfFIFI module when integrated into the excitation path of an iSIM^14,31^ (**Supplemental information, Supplemental Figure 4**). To visualize the excitation illumination of the iSIM, we used a concentrated dye solution^28^, and imaged the emitted light onto the camera without scanning (**Supplemental information, Supplemental Figure 4a**). Using an approximately Gaussian beam (M^2^<1.1, FWHM diameter ∼12 mm) to create the excitation pattern, as used in the initial iSIM design^9,29^, results in excitation points of spatially varying intensity closely following a Gaussian distribution (**Figure 2a**), as expected from the simulation (**Figure 1a**). Scanning the Gaussian excitation points partially homogenizes the excitation along the scan direction; however, the resulting illumination features a bright central region and strong roll-off away from the central optical axis (**Supplemental information, Supplemental Figure 5a,b**). In comparison, generating the excitation using the mfFIFI module results in a more uniform multifocal excitation covering an area of approximately 100×100 µm^2^ (**Figure 2b**), and a more homogeneous excitation during scanning (**Supplemental information, Supplemental Figure 5c-e**).

**Figure 2.**
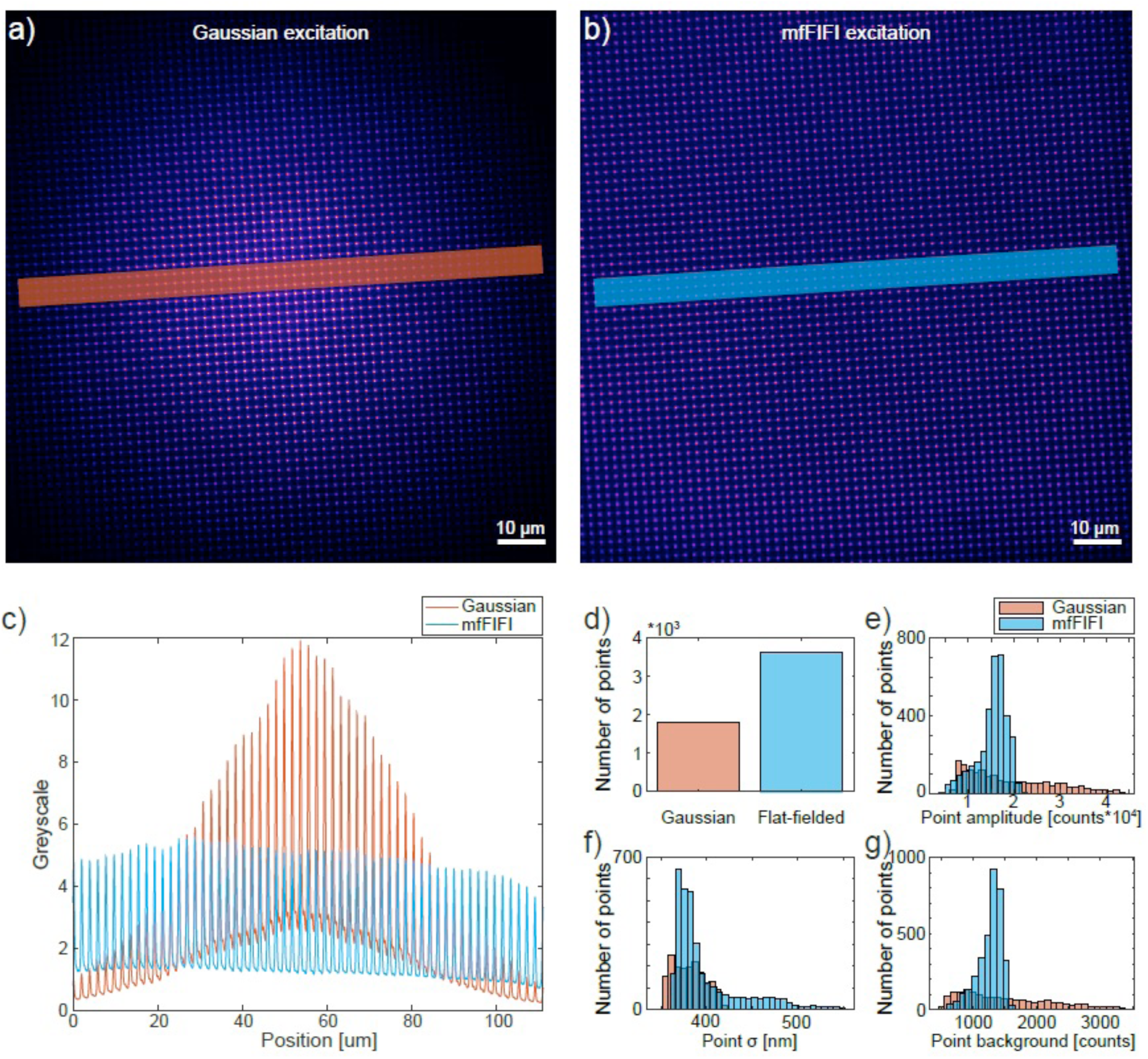
Performance comparison between Gaussian and mfFIFI excitation. a,b) Excitation points imaged onto a NaFITC fluorescent dye sample with (a) Gaussian and (b) mfFIFI excitation. c) Intensity profiles measured along lines in (a, b). d) Number of excitation points with a signal-to-noise ratio (SNR) above 3 in Gaussian and mfFIFI excitation. mfFIFI increases the number of excitation points by a factor of two. e-g) Histograms of excitation point (e) amplitudes, (f) variance and (g) background levels for Gaussian and mfFIFI excitation.

Comparing intensity profiles along the rows of excitation spots shows that mfFIFI efficiently redistributes the spatially varying input Gaussian excitation over a larger area (**Figure 2c**). Quantifying these differences, we find that mfFIFI doubles the number of excitation spots above background levels compared to the Gaussian excitation, while resulting in a much narrower distribution of spot amplitudes and background (**Figure 2d-e, g**). Furthermore, this improvement comes at no measurable cost to the quality of the structured illumination. We find that both the size of the excitation spots required for resolution improvement in confocal microscopy (**Figure 2f**) and their periodicity (222.3±0.3 µm (Gaussian) and 222.2±0.3 µm (mfFIFI) (mean ± S.D.)) are maintained, and compare well with the ground truth (222 µm excitation MLA pitch) (**Supplemental information**). This was further verified by imaging 100 nm fluorescent beads, showing comparable lateral resolution between mfFIFI (214 ± 5 nm, mean ± S.D.) and Gaussian illumination (219 ± 12 nm), based on FWHM measured on raw iSIM data before deconvolution, or ∼140 nm lateral and ∼350 nm axial resolution after deconvolution^9^ (**Supplemental information**).

### Large FOV iSIM imaging and FOV stitching

We used our custom-built iSIM setup as a platform to combine the capability of mfFIFI excitation with a scientific complementary metal-oxide semiconductor (sCMOS) camera for detection to perform large FOV volumetric imaging at doubled resolution. While existing commercial SIM setups are mostly limited to FOVs with a linear dimension of 30-60 µm, our setup reaches >100×100 µm^2^, thus providing a ∼4- to 10-fold increase in FOV area. We used mfFIFI iSIM to simultaneously image multiple mammalian cells within a single FOV at doubled resolution (**Figure 3a, Supplemental Movie 1**). Accessible imaging speeds are limited merely by the frame rate of the detector; thus, frame rates of up to 100 Hz as reported for iSIM^9,29^ are preserved, since the increase in FOV is achieved by improved parallelization from the increased number of illumination spots. Thus, the increase in FOV translates directly into an increase in throughput.

**Figure 3.**
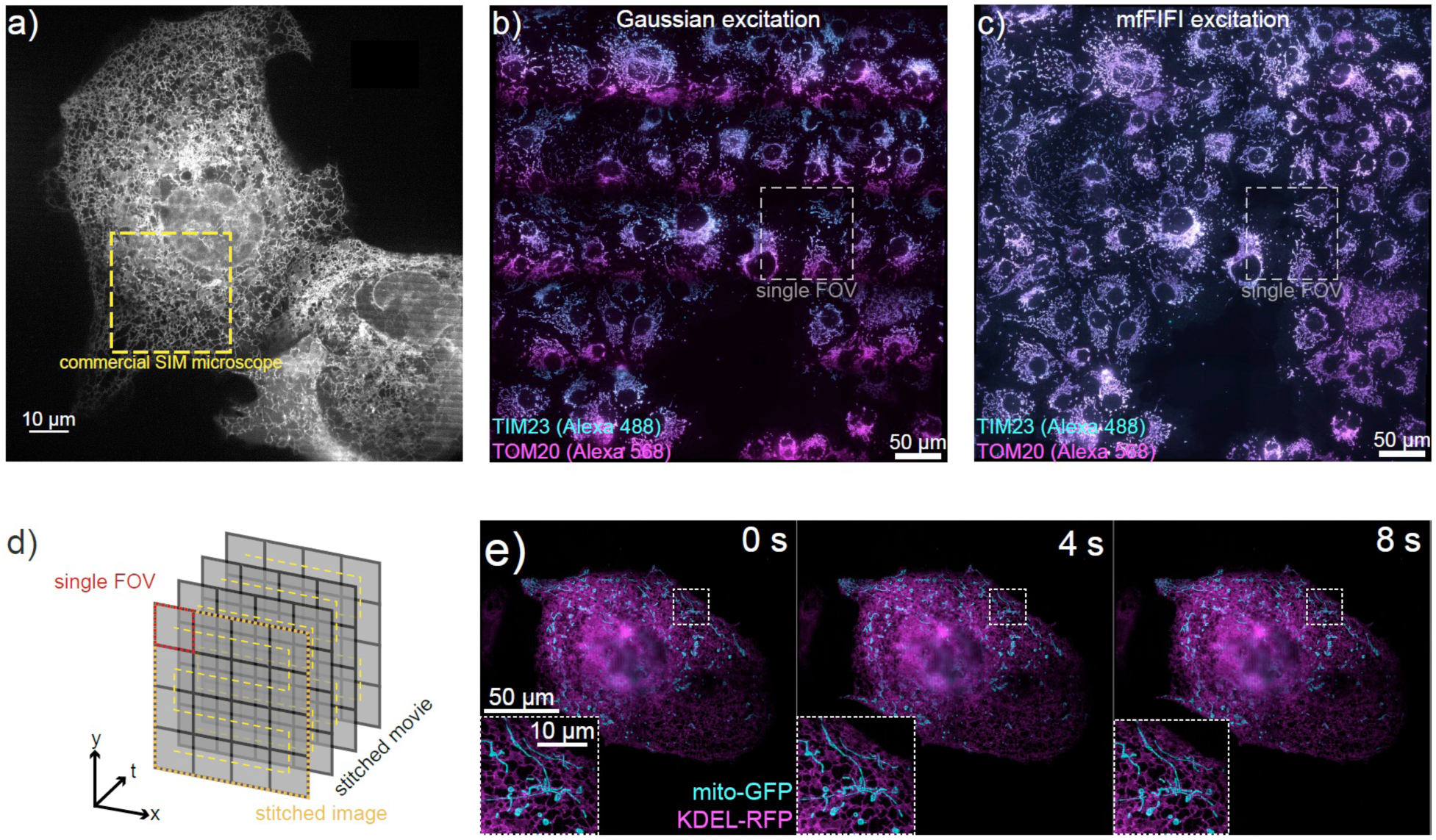
Large FOV imaging and multi-FOV stitching. a) Cos7 cell expressing KDEL-RFP imaged with an iSIM using mfFIFI excitation and an sCMOS camera enabling imaging of FOVs up to ∼115×115 µm^2^ at doubled resolution compared to diffraction limited performance. Yellow square shows the size of a 30×30 µm^2^ FOV available on a standard commercial SIM setup. Full movie shown in **Supplemental Movie 1**. b,c) Raw iSIM images of Cos7 cells immunolabelled for TIM23 and TOM20, stitched from a 5×5 grid of FOVs using (b) Gaussian and (c) mfFIFI excitation. d) Introducing an additional layer of parallelization into iSIM imaging by iterating over multiple FOVs allows trading off the speed of iSIM imaging to construct larger FOVs at high speed and doubled resolution compared to diffraction limited performance. e) Montage showing frames from a high-speed movie built up over a 2×2 grid of FOVs (**Supplemental information**, original movie 2s temporal resolution) of cells expressing mito-GFP and KDEL-RFP. Full movie shown in **Supplemental Movie 2**.

The effective image size can be further extended by stitching together adjacent FOVs. Such an approach requires homogeneous illumination to allow seamless stitching (**Figure 3b**). The speed of iSIM imaging combined with the large FOV enabled by mfFIFI allowed us to stitch together dual-color 3D stacks covering a 500×500×5 µm^3^ volume with a 5×5 grid of FOVs, each acquired within 5 seconds. With this approach, within 2 minutes, we could capture 3D images of more than 80 cells stained for the mitochondrial inner (TIM23) and outer (TOM20) membrane translocases, revealing coupled geometries of the two membranes (**Figure 3b-c**). The current limiting factor for multi-FOV imaging is the speed of stage translation and synchronization between stage translation and the rest of the iSIM acquisition control (**Supplemental information**). In contrast, the same acquisition without mfFIFI resulted in varying intensity across each FOV, with pronounced dark boundaries at the borders of individual FOVs (**Figure 3b**).

For time-lapse imaging, we introduced an additional layer of parallelization by iterating the acquisition procedure over a grid of FOVs (**Figure 3d**). By trading temporal resolution for a gain in imaging area, we can use the speed of iSIM imaging to cover multiple FOVs. For example, by iterating over a 2×2 grid of FOVs we could capture four time more regions of close contact between mitochondria and ER, known to serve as hotspots for mitochondrial division^30^ at a temporal resolution of 2 seconds (**Figure 3e, Supplemental Movie 2**) (**Supplemental information**). This illustrates how processes with less demanding temporal dynamics can benefit from mfFIFI to extend the throughput of iSIM without compromising spatial resolution.

### High-throughput super-resolution imaging of expanded centrioles

A recently developed variant of ExM termed ultrastructure expansion microscopy (U-ExM)^31^ enables preservation of the structure and molecular identity of multi-protein assemblies (particles). When combined with iSIM imaging, an expansion factor of ∼4 results in an effective resolution of ∼140 / 4 = 35 nm laterally and ∼350 / 4 ≈ 90 nm axially after deconvolution. Compared with current state-of-the-art super-resolution microscopes with similar resolution performance, such as the HT-STORM^10^ capable of imaging a 100×100 µm^2^ FOV in ∼5-10 minutes or a similar FOV with easySLM-STED^32^ in 60-80 minutes, the iSIM is capable of stitching together images of expanded samples to form an equivalent FOV within 2-5 seconds. This represents a 100-1000-fold improvement in imaging throughput. Therefore, acquiring datasets of thousands of particles that would take weeks or months to acquire on a conventional STORM or STED microscope require only 1-2 hours on the high-throughput iSIM.

As a proof-of-concept for iSIM/U-ExM, we set out to analyze centrioles, organelles found in most eukaryotic cells that seed the formation of the axoneme in cilia and the centrosome in animal cells. Centrioles have a characteristic nine-fold radial symmetric arrangement of microtubule triplets towards the proximal end of the organelle, composed of complete A-microtubules and incomplete B- and C-tubules^33^, with a transition to A- and B-microtubule doublets towards the centriole distal end^34^ (**Figure 4a**). It is known that centriolar tubulin is enriched in post-translational modifications (PTMs), including acetylation and polyglutamylation^35,36^. Acetylation is known to stabilize microtubules, increase their flexibility and protect them against mechanical aging^37–39^. Glutamylation has been postulated to likewise stabilize microtubules in the centriole and protect the organelle from pulling and pushing forces acting during mitosis^36,40,41^. Although these observations indicate that centrioles might be stabilized via tubulin acetylation and polyglutamylation, the spatial organization of these PTMs on the organelle remains largely unexplored.

**Figure 4.**
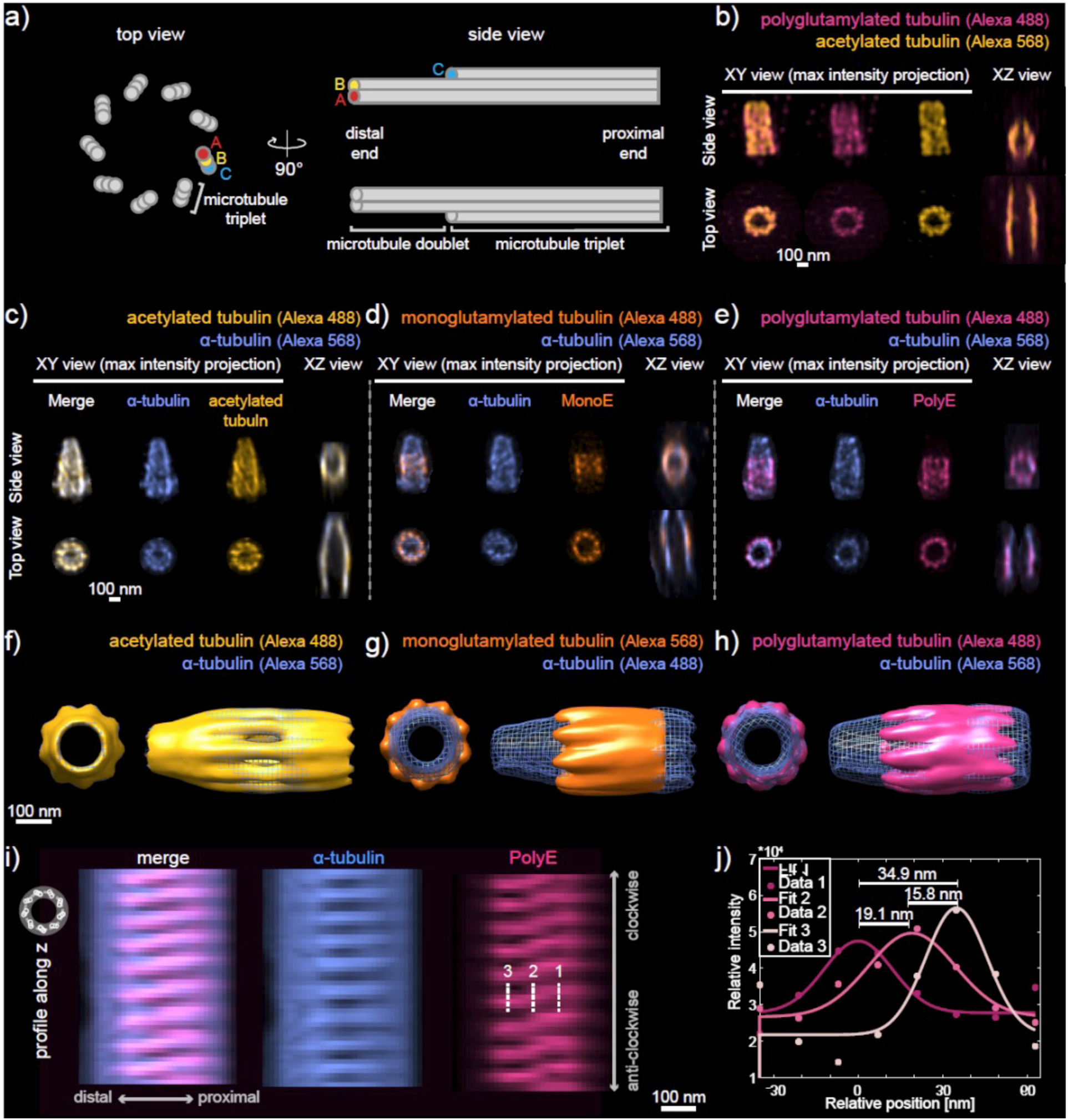
High-throughput super-resolution imaging of expanded centrioles for mapping post-translational modifications of centriolar tubulin. a) Schematic representation of the centriolar microtubule wall viewed from the side (left) and top from the distal end. The A-, B- and C-microtubules are indicated. b) Examples of individual mature human centriole particles orthogonal (side view) or parallel (top view) to the optical axis collected in expanded RPE-1 cells labelled for acetylated tubulin and polyglutamylation (PolyE). The lateral (XY) maximum intensity projection and axial (XZ) cross sections are shown. c-e) Examples of individual purified *Chlamydomonas reinhardtii* centriole particles viewed orthogonal (side view) or parallel (top view) to the optical axis labelled for α-tubulin as reference and (c) acetylated tubulin, (d) monoglutamylated tubulin (GT335, MonoE) and (e) polyglutamylated tubulin (PolyE). f-h) High resolution particle averaging reconstructions of (f) acetylated tubulin, (g) MonoE and (h) PolyE with α-tubulin as reference. Scale bars: 100 nm (pre-expansion). Top views shown from the distal end. i) Circular axial projection (XZ profile) of the centriole reconstruction from (h) showing α-tubulin and PolyE signal around the centriole (viewed from the outside). j) Intensity profiles measured along the dashed lines from (i) showing the radial displacement of the PolyE signal.

To analyze human centrioles in a similar stage of the cell cycle and of the centriole duplication cycle, we synchronized RPE-1 cells with thymidine to arrest them at the G1/S transition and with the Plk4 inhibitor Centrinone to prevent the formation of new centrioles^52^. After expansion and staining with antibodies against acetylated and polyglutamylated tubulin, we imaged >100 centrioles in 3D in their cellular context per hour, revealing the distribution of these PTMs along the organelle. Analyzing over 400 individual particles extracted from these images (**Supplemental Figure 6**) revealed that centriolar tubulin appears uniformly acetylated, while polyglutamylation terminates ∼30 nm before the distal end (**Figure 4b, Supplemental Figure 7a-c**). Moreover, the polyglutamylation signal exhibits a larger diameter (**Figure 4b, Supplemental Figure 7a-c**), in line with the fact that acetylation occurs witin the microtubule wall and polyglutamylation on the outer surface of the polymer^31^.

To further increase throughput, we sought to expand purified centrioles to capture dozens of pairs within a single FOV rather than just one per cell. We used centrioles purified from *Chlamydomonas reinhardtii* to this end since the U-ExM protocol had been optimized for this species^31^. Similarly to what we observed with human centrioles in the cellular context, we observe that in centrioles purified from *Chlamydomonas reinhardtii*, acetylated tubulin colocalizes with the microtubule wall along the entire length of the organelle (**Figure 4c**), in agreement with previous reports^42^. In contrast, we found that tubulin monoglutamylation (MonoE) is concentrated in the central core region of *Chlamydomonas reinhardtii* centrioles (**Figure 4d**), as previously reported^42^. Since MonoE recognizes the first glutamylation branching required for polyglutamylation (PolyE), we would expect the distribution of PolyE to partially or completely overlap with that of MonoE. Surprisingly, we found that although PolyE also localizes on the outer surface of the microtubule triplets where it has been suggested to localize to the C-microtubule^31^, it covers a wider band along the length axis than MonoE (**Figure 4e, Supplemental Figure 7, 12**). This difference could reflect lower MonoE antibody labelling efficiency, since saturating the signal shows similar MonoE/PolyE localization.

The high throughput offered by mfFIFI-iSIM allowed us to collect 3D images of thousands of purified expanded *Chlamydomonas reinhardtii* centrioles per hour (**Supplemental Figure 8, Supplemental Table 2**). Having obtained 3D rather than 2D particles allowed us to use fewer particles to perform averaging and high-resolution multicolor reconstruction^44^ compared to previous approaches using STORM^45^ or electron microscopy^46,47^. Thus, we classified particles before reconstruction to identify and average only those particles sharing a high degree of similarity (ranging from 10-50% of the whole dataset), and thereby achieve even higher resolution reconstructions (**Figure 4f-h, Supplemental information, Supplemental Figure 9**). Here, we have focused on the most frequently appearing class, but other classes could also be independently reconstructed. For mapping multiple PTMs within the centriole, we employ dual-color labeling, using α-tubulin staining of the microtubule triplet wall as a reference to subsequently align all PTMs (**Supplemental Figure 10**).

The resulting reconstructions revealed an unexpected shift in the tangential position of the PolyE signal from the proximal to the distal end (**Figure 4h**), which is also sometimes visible in individual raw particles (**Supplemental Figure 8b inset, Figure 4e**). To account for this shift, we considered two possibilities: 1) the whole microtubule triplet twists around its axis while the PolyE signal remains localized to the C-microtubule^48^ giving it an apparent twist, and 2) the PolyE signal changes microtubule localization within the microtubule triplet. Arguing against the first possibility, the microtubule triplet twists only within ∼100 nm of the proximal end (Le Guennec et al., in press), while in our reconstruction the PTM twist occurs within a wider region covering ∼250 nm from the proximal end,. Moreover, we determined that the first possibility would be expected to cause a moderate ∼6 nm shift of the PolyE signal, and the second a more pronounced ∼35-40 nm shift (**Supplemental information, Supplemental Figure 11**). From our reconstruction, we measure a twist of ∼34.9 nm, which cannot be explained purely by twisting of the microtubule triplet (**Figure 4i,j, Supplemental information, Supplemental Figure 11**). Therefore, we propose that polyglutamylation changes localization within the microtubule triplet along the centriole, moving from A-on the proximal end to the C-microtubule more distally.

## Conclusion

To summarize, we designed a flat-fielding method for efficient multifocal excitation, accompanied by a wave optics simulator which ensures optimal resolution performance. While the Koehler integrator has spot generating properties when used with coherent illumination, it introduces a strong wavelength dependence of the periodicity, making it unsuitable as such for multi-color applications^26^. mfFIFI provides a largely wavelength-independent solution for generating homogeneous multi-focal excitation, while benefiting from the homogenizing properties of the Koehler integrator. We implemented the mfFIFI module into an iSIM microscope, since its fast acquisition speeds and resolution doubling give it good potential for high-throughput super-resolution imaging. However, the mfFIFI module could be extended to any multi-focal excitation microscope such as a spinning disk confocal microscope^6^ using a Nipkow disk configuration, but not with other flat-fielding solutions using diffractive optical elements^20^, spatial light modulators (SLM)^21^ or beam splitters^23^. Furthermore, combining mfFIFI with phase masks required for donut beam shaping could extend its application to parallelized STED and RESOLFT microscopes.

Flat-fielding enables multi-field of view imaging, which is a powerful way to maintain resolution while increasing the effective image size to span more length scales. This is important for sample spanning multiple fields of view, as in the case of tissues, or expanded samples. ExM sacrifices the effective FOV to increase resolution, and stitching provides a way to maintain throughput, allowing cellular features or structures to be imaged *in situ*, as we demonstrate by imaging hundreds of human centrioles in cells.

Particle averaging and reconstruction is traditionally used in ultrastructural biology, and leverages large datasets to capture the intrinsic variability within particles through classification before reconstruction, while averaging out the noise. In the case of ExM, different gels can have slightly different expansion factors, but this issue is circumvented by our iSIM setup since thousands of particles can be acquired from a single gel section (∼5×5 mm^2^). In our case, the collected datasets were sufficiently large to allow particle classification prior to averaging, an approach which improves the resolution of the reconstruction and which, to our knowledge, has not been previously used for fluorescence super-resolution. In addition to a 100-1000-fold improvement in throughput of iSIM/U-ExM over HT-STORM^10,45^ or easySLM-STED^32^, expanded samples offer enhanced accessibility of antibodies to bind proteins in crowded assemblies. Beyond particle averaging, high-content or machine learning approaches rely on large datasets for training or screening, and can benefit from the combination of throughput and resolution described here.

Our previous attempts to image centriolar microtubules with STORM proved largely unsuccessful, restricting previous studies to the more accessible epitopes^45^. Here, we reveal previously unobserved molecular organization, showing high coverage of polyglutamylation in human centrioles, as well as precise localization to microtubule triplets along the *Chlamydomonas reinhardtii* centriole. We find that polyglutamylation appears to shift from the A-microtubule proximally to the C-microtubule more distally. It is possible that such a glutamylation shift reflects a centriolar tubule code that defines the spatial boundaries of the various elements that compose it, which could determine the positioning of different structural elements along the length of the organelle.

## Methods

### Sample preparation

Cos-7 and RPE-1 cells were grown in Dulbecco’s modified Eagle medium (DMEM) supplemented with 10% fetal bovine serum (FBS). Cells were plated on 25 mm, #1.5 glass coverslips (Menzel) 16-24 h prior to transfection or fixation at a confluency of ∼10^5^ cells per well.

Transfections using [insert], were performed with Lipofectamine 2000 (Life Technologies) using 150 ng of [insert] and 1.5 µL of the Lipofectamine 2000 reagent in 100 µL Opti-MEM medium per 6 well.

For synchronization of RPE-1 cells at the G1/S transition and the onset of centriole assembly, ∼125’000 cells were seeded for 24 h in six well-plates (Merck, TPP, 92006) on 12 mm coverslips in DMEM [insert]. Thymidine (Merck, T1895, 1 mM) and Centrinone (Lucerna-Chem, MCE-HY-18682, 300 nM) were added for 18 h, before fixation of cells with -20°C methanol for 5’.

For immunofluorescence experiments with COS-7 cells, cells were washed in pre-warmed PBS before being fixed in pre-warmed fixation buffer (4% paraformaldehyde in PBS). Cells were then permeabilized in 0.25 % Triton-X in PBS for 10 min. After washing in PBS for 5 min, cells were incubated in blocking buffer (1% BSA in PBS) for 60 min. The primary antibodies (Tom20-rabbit (1:50, FL-145 sc-11415 Santa Cruz Biotechnologies), Tim23-mouse (1:100, 611222 BD Biosciences)) diluted in blocking buffer were incubated for 60 min before washing 3 times in 0.2% BSA with 0.25% Triton-X in PBS for 10 min. Secondary antibodies (AlexaFluor 488 goat anti-mouse IgG (H+L) (1:150, A28175 ThermoFisher), AlexaFluor 568 goat anti-rabbit IgG (H+L) (1:150, A11011 ThermoFisher)) were diluted in blocking buffer before incubation for another 30 min. The sample was incubated in the dark, then washed 3 times with PBS before imaging.

### iSIM imaging

The iSIM setup was partly based on previously described implementations^9,29^. Two lasers with wavelengths of 488 nm (Sapphire 488-300 CW CDRH, Coherent) and 561 nm (gem 561, Laser Quantum GmbH) were combined using a dichroic mirror (F48-486, Analysentechnik) and controlled through an acousto-optic tunable filter (AOTFnC-400.650-TN, AA Optoelectronic). In the case of Gaussian excitation, the beam was expanded with a 10× beam expander (f_1a_ = 40 mm, f_2a_ = 400 mm, Thorlabs). In the mfFIFI path, a focusing lens (f_FC_ = 50 mm) was used to focus the light near a rotating diffuser (2.5° ± 0.25° FWHM at 650 nm, 24-00066, Süss MicroOptics SA) before collimation by a collimating lens (f_CL_ = 60 mm, Thorlabs). The collimated beam was contracted by a factor of 4 by two lenses (f_1b_ = 120 mm, f_2b_ = 30 mm, Thorlabs) before illuminating two flat-fielding MLAs (300 µm pitch, 10mm × 10 mm, f = 4.78 mm, square lenses, 18-00157, Süss MicroOptics SA). The flat field was then focused by a Fourier lens (f_FL_ = 300 mm, Thorlabs). Both paths then illuminated the excitation MLA (222 µm pitch, 1” diameter, f = 6 mm, square lenses APO-Q-P222-R2.74, Advanced Microoptic Systems GmbH). The excitation was relayed by scan lenses (f_SL_ = 190 mm, 55-S190-60-VIS, Special Optics) to and from the scanning mirror (SPO9086 Rev B Coated X Mirror, Sierra Precision Optics) mounted on a galvanometer scanner (QS-12, N-2071, Nutfield Technology). The excitation was then imaged onto the sample using a tube lens (f_TL_ = 350 mm, 49-289-INK, Edmund Optics) and an objective lens (APON60XOTIRF, Olympus), resulting in a final magnification of ∼116x. The sample was placed on a precision aligned microscopy platform including a micropositioning control (RM21-AZ-AXY-RMS-M, Mad City Labs) and a Z piezo stage (Nano-Z200, Mad City Labs). The emission was relayed back through the excitation side of the scanning mirror and split using a dichroic mirror (F58-488S, Semrock). The emission was then pinholed with a pinhole array (chrome on 0.090-inch-thick quartz, 222 µm pitch, 40 µm diameter, Photosciences) and relayed by two relay lenses (f_R_ = 300 mm, Thorlabs) before contracting each emission spot by a factor of 2 using another MLA (222 µm pitch, 1” diameter, f = 0.93 mm, square lenses APO-Q-P222-R0.425, Advanced Microoptic Systems GmbH). The emission was then relayed using scan lenses on the emission side of the scan mirror, towards a filter wheel (Lambda 10-B, Sutter Instruments, Science Products) with 2 notch filters (NF-03-488E-25 and NF-03-561E-25, Semrock) mounted to block the excitation wavelengths. The transmitted emission was then collected by a sCMOS camera (PrimeBSI, 01-PRIME-BSI-R-M-16-C, Photometrics). The size of a square camera pixel corresponds to 56 nm on the sample.

The microscope was controlled using a custom written MATLAB script which controlled an analog output card (PCI-6733, National Instruments) and breakout box (BNC-2110, National Instruments) for precise control and synchronization of the scan mirror, AOTF, camera and Z-stage. The XY stage and camera were controlled through MicroManager^49^.

For live cell imaging, imaging was performed at 37 °C in pre-warmed Leibovitz medium in a top stage incubator (H301-PRIOR-NZ100-H117, Okolab). Imaging of immunostained samples was performed in PBS. Expanded samples were mounted as previously described^31^. Briefly, the expansion factor was determined before imaging by measuring the gel size with a caliper (precision ±0.01 mm). The gel was then cut using a razor, before removing excess water and placing the gel on a poly-D-lysine treated coverslip (25 mm round coverslips, Mendel #1.5) already placed inside the imaging chamber. After gently pressing on the gel to ensure attachment to the coverslip, a few drops of ddH_2_O were added on top of the sample to avoid shrinkage during imaging. Image acquisition was performed using a custom written MATLAB script, in combination with MicroManager^49^.

### iSIM deconvolution

Raw iSIM images were deconvolved using the Lucy-Richardson deconvolution algorithm implemented in MATLAB and provided by Dr. Hari Shroff^9^. Each raw z-stack was deconvolved for 40 iterations.

### mfFIFI characterization platform

An optical characterization setup was built to measure the variation in pitch and spot size of the multifocal points. A laser beam with a wavelength of 640 nm (CUBE 640-100C, Coherent) was expanded with a 6x beam expander (f = 50 mm, f = 300 mm, Thorlabs) with a pinhole (P50D, Thorlabs) at the joint focal plane of the two lenses to spatially filter the beam. The beam is passed through a Köhler integrator, made of the following components: focusing lens (f = 80 mm, Thorlabs), rotating diffuser (2.5° ± 0.25° FWHM at 650 nm, 24-00066, Süss MicroOptics SA), collimating lens (f = 40 mm, Thorlabs), two flat-fielding MLAs (300 µm pitch, 10mm × 10 mm, f = 4.78 mm, square lenses, 18-00157, Süss MicroOptics SA), Fourier lens (f_FL_ = 300 mm, Thorlabs). An iris (ID25, Thorlabs) is used before the two MLAs to control the portion of the beam which is allowed to propagate forward to reach and illuminate MLAs, by adjusting the opening diameter of the iris. An excitation MLA (300 µm pitch, 10mm × 10 mm, f = 8.72 mm, square lenses, 18-00221, Süss MicroOptics SA) is used to produce the multifocal points. The size of the multifocal illumination is reduced by a factor of 0.6x by a beam de-expander (f = 50 mm, f = 30 mm, Thorlabs) to be of appropriate size for the sensor of the camera (DCC1545M, Thorlabs).

To obtain the data shown in Figure 1e, h, we used the characterization platform to test the parameters affecting the spot size and pitch. A Vernier caliper was used to measure the iris opening diameter, which was varied methodically to control the size of the beam which illuminates the MLAs. Images of the beam at different iris opening diameters were analyzed using a MATLAB script, which allowed us to fit each excitation point to a 2D Gaussian and measure the multifocal points’ mean spot FWHM as function of iris diameter. In order to study the variation in the pitch of the multifocal points, the distance between the Fourier lens and the second flat-fielding MLA was varied by displacing the flat-fielding MLAs. A MATLAB script is used to measure the mean pitch at each distance, so that the relationship between this distance and the pitch can be determined.

### Image analysis

#### Plateau uniformity quantification

The homogeneity was quantified using the plateau uniformity definition based on the FWHM of the histogram of the intensity values, according to ISO, ISO 13694:2018: Optics and optical instruments - lasers and laser-related equipment - test methods for laser beam power (energy) density distribution.

#### Illumination profile measurement

The illumination profiles used to optimize the pitch (telecentricity) and spot size of the multifocal excitation were visualized using a small color CMOS camera (DCC1645C, Thorlabs), placed in an intermediate image plane of the iSIM, between the second scan lens and tube lens, to avoid introducing other aberrations along the whole optical path of the iSIM.

The excitation profile at the sample was measured using a highly concentrated fluorescent dye sample^32^, as previously described^9^. Briefly, sodium fluorescein (NaFITC) powder was diluted in deionized water, followed by vortexing and sonication until the powder was completely dissolved. A 10 µL drop was placed on a 25 mm coverslip, before covering with a 12 mm coverslip. The sample was then sealed using nail polish. The same dye sample was used to measure both the 488 nm and 561 nm illumination profiles. The illumination spots were characterized by fitting a 2D Gaussian to the intensity image. The threshold was set so that no points in the background were detected. The scanning profiles were measured with a 150 pixel thick line in Fiji^50^.

In the case of excitation spots visualized in Figure 1f, g and the quantification in Figure 1e, h, the illumination profile was imaged directly onto a camera placed in a conjugate image plane.

#### Bead FWHM analysis

To characterize the performance of the iSIM microscope, we used 100 nm Tetraspeck beads (ThermoFisher Scientific, T7279) deposited on poly-L-lysine treated coverslips. Bead size was determined by fitting a Gaussian profile to the intensity profile of the bead and extracting the FWHM.

#### Centriole shape and coverage analysis

To analyze the diameters of different PTM localizations within the centrioles, we measured the intensity profile through the centers of top view centrioles (central plane, centriole barrel parallel to the imaging axis). The profile was measured using a 7 pixel thick line in Fiji^50^. Each peak was fitted to a Gaussian profile to localize its center, before calculating the diameter as the distance between the two peaks on opposing sides of the centriole ring.

To analyze the PTM distribution along the length of the centriole and their coverage, we measured the intensity profile along the length of the centriole using a 10 pixel thick line and from summed intensity projections of side view centrioles (centriole barrel orthogonal to the imaging axis). We then took the width of the profile at ¼ of the maximal signal as the length, to be less susceptible to noise. The coverage was then calculated by dividing the length of the PTM in question, by the length of the reference label (acetylated tubulin for human centrioles and α-tubulin in the case of *Chlamydomonas reinhardtii* centrioles).

#### Multiple field of view stitching

For stitching of multiple images, raw images or z-stacks were acquired with a 10% overlap for both Gaussian and mfFIFI excitation. The acquired images were then combined using the Grid/Collection stitching tool in Fiji^50^ using the linear blending method with the default values of 0.30 for regression threshold, 2.50 for max/avg displacement threshold and 3.50 for absolute displacement threshold.

#### Wave optics simulations

The mfFIFI simulation platform has been adapted based on previous work^9^. All simulations are based on standard numerical Fourier optics algorithms and rely on angular spectrum propagation methods to simulate the propagation of an optical wave. An initial simulation showing the effect of illuminating the excitation MLA with Gaussian and flat-fielded profiles was performed using phase masks to represent optical components. The parameters used for the simulation are summarized in **Supplemental Table 1**.

#### Centriole expansion protocol

*Chlamydomonas reinhardtii* centrioles were isolated from the cell-wall-less strain CW15-^51^ and expanded using the U-ExM protocol^31^. Briefly, isolated centrioles were spun on 12mm Poly-D-Lysine-coated (ThermoFisher, A3890401) coverslips prior to U-ExM. For human cell expansion, cells were initially seeded on 12 mm coverslips before being fixed with 4% PFA. Coverslips were then incubated in a solution of 0.7% formaldehyde and 1% acrylamide in PBS for 4-5 hours at 37°C. Next, coverslips were incubated in the monomer solution for 1 minute on ice and then shifted to 37°C, for 1 hour in a dark and humidified chamber. For one gel, the monomer solution is made of 25 µL sodium acrylate (Sigma-Aldrich, 408220, 38% (wt/wt, diluted with nuclease-free water)), 12.5 µL acrylamide (Sigma-Aldrich, A4058, 40% stock solution), 2.5 µL N,N’-methylenbisacrylamide (Sigma-Aldrich, M1533, 2% stock solution), and 5 µL 10X PBS, supplemented with 2.5 μL TEMED and 2.5 μL APS (from 10% stock solutions). Once polymerized, gels were moved into denaturation buffer (200 mM SDS, 200 mM NaCl and 50 mM Tris in nuclease-free water, pH 9) for 15 minutes at room temperature with gentle shaking and then shifted for 30 minutes to 95°C in an 1.5 mL Eppendorf tube with 1 mL of fresh denaturation buffer. Then, gels were placed at room temperature in beakers with 200 mL of distilled water. Water was exchanged twice (every 30 min) and the sample was incubated overnight at room temperature. The following day, water was changed with PBS for 15 min at room temperature. Gels were then incubated for 3h at 37°C with gentle shaking in 1 mL of primary antibody solution (in 3% BSA and 0.05% Tween20). Samples were then washed three times for 10 min with PBS supplemented by 0.1% Tween20 while shaking, followed by incubation with secondary antibodies for 3h at 37°C in 3% BSA and 0.05% Tween20. Finally gels were washed 3x in T-PBS. For RPE-1 cells, the sample was supplemented in the second wash with Hoechst 33258 (1:2’000) dye and place in beakers with 200 mL of distilled water for final expansion, with again water being exchanged twice for 30’. Before imaging, gels were again expanded overnight in ddH_2_0. Primary antibodies used to image centrioles in expanded RPE-1 cells were rabbit anti-polyglutamate chain (polyE), pAb (IN105, 1:250) and mouse monoclonal anti-acetylated tubulin (Merck, T7451, 1:500). Secondary antibodies were goat anti-Rabbit Alexa488 (Thermo-Fisher Scientific, A11034, 1:500) and goat anti-mouse IgG Alexa 568 F(ab’)2 (Thermo-Fisher Scientific, A11019, 1:500).The following primary antibodies were used for imaging expanded purified *Chlamydomonas reinhardtii* centrioles: rabbit polyclonal anti-polyglutamate chain (PolyE, IN105,1:500, AG-25B-0030-C050, Adipogen), mouse monoclonal anti-α-tubulin (DM1α,1:500, T6199, Sigma-Aldrich), mouse monoclonal anti-polyglutamylation modification (GT335,1:200, AG-20B-0020, Adipogen), mouse monoclonal anti-acetyl-α-tubulin (Lys40,1:50, 32-2700, Invitrogen, ThermoFisher). Secondary antibodies were goat anti-rabbit Alexa Fluor 488 IgG H+L (1:400, A11008), goat anti-mouse Alexa Fluor 488 IgG H+L (1:400, A11029), goat anti-rabbit Alexa Fluor 568 IgG H+L (1:400, A11036), goat anti-mouse Alexa Fluor 568 IgG H+L (1:400, A11004 all from Invitrogen, ThermoFisher).

#### Particle segmentation, classification and 3D reconstruction of purified centrioles

Particles were segmented using a custom written Matlab script. Briefly, the threshold value was found on the maximum intensity projections of the 3D stacks, before applying it back to the individual stacks. The regions were then binarized and segmented by connecting neighboring pixels across the 3D stack. The binarized points of interested were then dilated to form the mask which was then applied back to the raw stack. The particles were then cut out from the original stack with their specific mask and a 60×60 pixel surrounding region. The segmentation was ∼50% successful, mostly because centrioles are often found in pairs, and closely located pairs had to be rejected since they could not be segmented correctly.

After particles were segmented and up-sampled (to reach isotropic pixel size), the particles with tubulin signal were aligned to a reference using Dynamo. The reference was built using a reconstructed tubulin signal made from 12 manually selected particles. Once particles were all similarly aligned, a cross-correlation matrix was generated by calculating the similarity between each pairs of particles. The cross-correlation matrix was converted to a distance matrix by subtracting cross-correlation values to 1. A hierarchical classification was made using the Ward method (hclust function from R). The classification tree was empirically cut into 10 groups. For each group, the average particle was generated for the tubulin signal and for the PolyE signal by applying the transformation parameters obtained during the alignment step. The 10 averages were compared to identify which groups were most promising. The best classes were used to produce the reconstruction results of Figure 4. The reconstruction method was based on ref.^44^, and followed the procedure reported in ref.^31^ with a C9 symmetry constraint. The point spread function was experimentally measured by imaging 100 nm fluorescent beads (TetraSpeck^TM^ Microspheres, 0.1 µm, T7279 ThermoFisher). We made two modifications to the *reference-free* step of the reconstruction algorithm^44^. Firstly, we adopted a multiscale reconstruction approach: we used results obtained with coarsely subsampled data to get a coarse initialization. Thanks to this approach, we were able to perform the reconstruction at the highest resolution in reasonable computation time. Secondly, we decoupled the angular search of the orientations: the first two Euler angles were estimated first, before the third one. This further accelerated the reconstruction and provided sharper results in the case of C9 symmetry.

Reconstructions were visualized using Chimera, by setting the threshold to fit the average length of the particles. Artefactual signals in the center of the reconstruction that arise due to the imposed 9-fold symmetry were removed during post-processing.

## Acknowledgements

We thank Hari Shroff and Alistair Curd for their help and advice for the construction of the iSIM; Cordelia Berz for her help with multi-FOV imaging; Raoul Kirchner on discussions on the Koehler integrator; Christian Sieben and Lina Carlini for critical reading of the manuscript.

This work is supported by the European Research Council (ERC; AdG 835322, CENGIN to P. Gö and ERC; StG 715289 ACCENT to P.Gu), the Swiss National Science Foundation (SNSF) PP00P3_157517 to P. Gu and 182429 to S.M., the MSCA (75200, CARTASSY to N.B.), and the National Centre for Competence in Research (NCCR) Chemical Biology (S.M.).

## Author contributions

D.M., K.M.D., P.Gö., V.H., P.Gu. and S.M. conceived and designed the project. P.Gö., V.H., P.Gu and S.M. supervised the project. D.M., K.M.D. and S.M. designed the multi-focal illumination system. D.M. and K.M.D. developed the simulation platform. D.M. built the microscope, performed all experiments and data analysis. D.G. performed the purified *Chlamydomonas reinhardtii* centriole sample preparation. D.F. performed single particle averaging and reconstruction. N.B. performed the human RPE-1 centriole sample preparation. M.L.G. performed the particle classification and alignment. K.I. built and performed experiments on the flat-fielding characterization platform. D.M. and S.M. wrote the manuscript with contributions from all authors.

## Competing interests

The authors declare no competing financial interests.

