## Supplemental Information for "Homogeneous multifocal excitation for high-throughput super-resolution imaging"

#### Design equations

Based on the requirements for homogeneous, structured multifocal excitation, we revise the Koehler integrator and propose the following design equations for its implementation in multi-focal confocal microscopes:

Fresnel Number[25,26]:

$$FN = \frac{p^2}{4\lambda f}$$

For good homogeneity FN≳5

Flat field size[25,26] (at excitation microlens array):

$$S = \frac{F_{FL} p}{f}$$

Depends on size of the field of view and magnification

Flat field size (at sample):

$$S_{sample} = \frac{1}{M} \frac{F_{FL} p}{f}$$

Size of excitation spots (in focal plane of excitation microlens array):

$$r = \frac{f \cdot f_x \cdot F_1}{F_{FL} \cdot F_{CL} \cdot F_2} R_{source} + \frac{f_x \cdot F_2}{F_{FL} \cdot F_1} R$$

Depends on magnification

Size of excitation spots (at sample) should be diffraction-limited:

$$r_{sample} = \frac{1}{M} \left( \frac{f \cdot f_x \cdot F_1}{F_{FL} \cdot F_{CL} \cdot F_2} R_{source} + \frac{f_x \cdot F_2}{F_{FL} \cdot F_1} R \right)$$

Flat field homogeneity:

$$B = \frac{R}{p}$$

For good homogeneity, B≳5

No crosstalk condition[26]:

$$\frac{F_1 \cdot f}{F_2 \cdot F_{CL}} \cdot R_{source} \leq \frac{p}{2}$$

The design equations are functions of the following design parameters:

$f$ – focal length of flat-fielding microlens arrays

$p$ – pitch of flat-fielding microlens arrays

$f_x$ – focal length of excitation microlens array

$p_x$ – pitch of excitation microlens array

$F_{FL}$ – focal length of Fourier lens

$F_{CL}$ – focal length of collimating lens

$F_1$ – focal length of first beam expander lens

$F_2$ – focal length of second beam expander lens

$R_{source}$ – radius of extended source

$\theta_{source}$ – angular divergence of extended source

$M$ – magnification

$R$ – beam radius incident on the first flat-fielding MLA without the beam expander

Can be estimated as: $R \cong R_{source} + F_{CL}\tan(\theta_{source})$ (see Ray Transfer Matrix Calculation)

$\lambda$ – wavelength

Furthermore, in the case of laser illumination, the size and the divergence of the extended source can be controlled by displacing the rotating diffuser, or changing the focal length of the focusing lens respectively. These design equations help identify the components necessary to implement the adapted Koehler integrator to multi-focal confocal systems. To our knowledge, the proposed illumination system should be easily adaptable to any multi-focal confocal microscope.

**Ray transfer matrix calculation**

To better understand the adapted illumination system, we performed a ray transfer matrix calculation for the whole system, as adapted from[27]. To fully explore the effect of introducing a beam expander between the collimating lens and the first flat-fielding microlens array, let us compare the two cases side-by-side. For this, we divide the system into three parts:

Sub-system 1: $(y_1, \beta_1) \rightarrow (y_2, \beta_2)$

Starting at the extended source

Ending in plane containing the first flat-fielding microlens array, but without considering its effect

$y_2$ effectively determines the radius of the beam incident on the first flat-fielding microlens array ($R$)

Without beam expander:

$$\begin{pmatrix} y_2 \\ \beta_2 \end{pmatrix} = \begin{pmatrix} \left(1 - \frac{L}{F_{CL}}\right) y_1 + F_{CL}\beta_1 \\ -\frac{y_1}{F_{CL}} \end{pmatrix}$$

Where $L$ is the variable distance between the collimating lens and the first flat-fielding microlens array.

With beam expander:

$$\begin{pmatrix} y_2 \\ \beta_2 \end{pmatrix} = \begin{pmatrix} \frac{1}{F_{CL}\cdot F_1\cdot F_2}\left(F_2^2(L_1-F_{CL}-F_1)+F_1^2(L_2-F_2)\right)y_1 - \frac{F_2}{F_1}F_{CL}\beta_1 \\ \frac{F_1}{F_2}\frac{y_1}{F_{CL}} \end{pmatrix}$$

Where $L_1$ and $L_2$ are the distances between the collimating lens and the first lens of the beam expander, and the distance between the second lens of the beam expander and the first flat-fielding microlens array.

Sub-system 2: $(x_1,\alpha_1) \rightarrow (x_2,\alpha_2)$

Starting in local coordinate system of a single lenslet of the first flat-fielding microlens array

Ending in the focal plane of the Fourier lens, containing the excitation microlens array, but without considering its effect

$$\begin{pmatrix} x_2 \\ \alpha_2 \end{pmatrix} = \begin{pmatrix} -\frac{F_{FL}}{f}x_1 \\ -\frac{1}{F_{FL}}(\alpha_1 f - np) \end{pmatrix}$$

Not explicitly modified by introduction of beam expander

Sub-system 3: $(z_1,\gamma_1) \rightarrow (z_2,\gamma_2)$

Starting in local coordinate system of a single lenslet of the excitation microlens array

Ends in local focal plane of a single lenslet of the excitation microlens array

$$\begin{pmatrix} z_2 \\ \gamma_2 \end{pmatrix} = \begin{pmatrix} f_x\gamma_1 \\ \gamma_1 - \frac{z_1}{f_x} \end{pmatrix}$$

Not explicitly modified by introduction of beam expander

These three subsystems are linked together by boundary conditions that transition from the local coordinate systems of microlens arrays to the global coordinate system. The boundary conditions are:

Boundary condition $(y_2, \beta_2) \rightarrow (x_1, \alpha_1)$: from global coordinate system to the local coordinate system of the flat-fielding microlens arrays

$$\begin{pmatrix} x_1 \\ \alpha_1 \end{pmatrix} = \begin{pmatrix} y_2 - np \\ \beta_2 \end{pmatrix}$$

Boundary condition $(x_2, \alpha_2) \rightarrow (z_1, \gamma_1)$: from global coordinate system to the local coordinate system of the excitation microlens array

$$\begin{pmatrix} z_1 \\ \gamma_1 \end{pmatrix} = \begin{pmatrix} x_2 - mp_x \\ \alpha_2 \end{pmatrix}$$

Regardless of the elements before the first microlens array, the ray-tracing matrix between the first microlens array and the focal plane of the excitation microlens array is given by (adapted from [30]):

$$\begin{pmatrix} z_2 \\ \gamma_2 \end{pmatrix} = \begin{pmatrix} -\frac{f_x}{F_{FL}}(\beta_2 f - np) \\ \frac{F_{FL}}{f_x \cdot f} y_2 - \frac{f}{F_{FL}}\beta_2 + \left(\frac{1}{F_{FL}} - \frac{F_{FL}}{f \cdot f_x}\right) np + \frac{mp_x}{f_x} \end{pmatrix} \cong \begin{pmatrix} -\frac{f_x}{F_{FL}}(\beta_2 f - y_2) \\ -\frac{f}{F_{FL}}\beta_2 + \frac{1}{F_{FL}} y_2 + \frac{mp_x}{f_x} \end{pmatrix}$$

Where $np$ indicates the radius of the beam incident on the first microlens array, equally corresponding to the maximal height of a beam incident on the array, $R = \max(y_2)$. Similarly, the maximal values of $y_1, \beta_1$ are set by the radius and angular divergence of the extended source:

$$\max(y_1) = R_{source}$$

$$\max(\beta_1) = \theta_{source}$$

By replacing the values of $y_2$ and $\beta_2$ into the system with or without a beam expander, we see that the overall size of the excitation spots can be minimized more effectively in the case including the beam expander. To find the size of the excitation spot $r$, we simply compute the maximum of $z_2, r = \max(z_2)$:

Without beam expander

$$r \cong \frac{f_x}{F_{FL}}\left(1-\frac{L}{F_{CL}}+\frac{f}{F_{CL}}\right)R_{source}+\frac{f \cdot F_{CL}}{F_{FL}}\theta_{source}$$

Where $L$ is the distance between the collimating lens and the first flat-fielding microlens array. We can identify a few solutions that simplify the solution and facilitate the contraction of the beam expander, but are not obligatory:

If $L = F_{CL}$

Beam height: $y_2 = F_{CL}\beta_1$ set purely by divergence of extended source

Beam radius: $R = F_{CL}\theta_{source}$ set purely by divergence of extended source

Spot size: $r \cong \frac{f_x \cdot f}{F_{FL} \cdot F_{CL}} R_{source} + \frac{f \cdot F_{CL}}{F_{FL}}\theta_{source}$

If $L = F_{CL} - f$

Beam height: $y_2 = \frac{f}{F_{CL}} y_1 + F_{CL}\beta_1$

Beam radius: $R = \frac{f}{F_{CL}} R_{source} + F_{CL}\theta_{source}$

Spot size: $r \cong \frac{f \cdot F_{CL}}{F_{FL}}\theta_{source}$ set purely by divergence of extended source

With beam expander

$$r \cong \frac{f_x}{F_{FL} \cdot F_{CL}}\left(\frac{F_2}{F_1}(L_1 - F_{CL} - F_1) + \frac{F_1}{F_2}(L_2 - F_2) - \frac{f \cdot F_1}{F_2}\right)R_{source} + \frac{f \cdot F_{CL} \cdot F_2}{F_{FL} \cdot F_1}\theta_{source}$$

Where $L_1$ is the distance between the collimating lens and the first beam expander lens, and $L_2$ the distance between the second beam expander lens and the first flat-fielding microlens array. Since the second term of the expression for r dominates, we can see that introducing a beam expander rescales the size of the excitation foci by a factor of F_2/F_1. In the case where F_2<F_1, this leads to a decrease in size. $L_1$ and $L_2$ can be modified allow flexible tuning of the radius $R$. We identify a couple of solutions which simplify the final form and facilitate the contraction by the beam expander but are not obligatory:

If $L_1 = F_1 + F_{CL}$ and $L_2 = F_2$:

Beam height: $y_2 = -\frac{F_2}{F_1} F_{CL}\theta_1$ set purely by divergence of extended source

Beam radius: $R^* = -\frac{F_2}{F_1} F_{CL}\theta_{source}$ set purely by divergence of extended source

Spot size: $r = \frac{f_X \cdot F_1 \cdot f}{F_{FL} \cdot F_{CL} \cdot F_2} R_{source} + \frac{f \cdot F_{CL} \cdot F_2}{F_{FL} \cdot F_1} \theta_{source}$

If $L_1 = F_1$ and $L_2 = F_2$:

Beam height: $y_2 = -\frac{F_2}{F_1}(y_1 + F_{CL}\beta_1)$

Beam radius: $R^* = -\frac{F_2}{F_1}(R_{source} + F_{CL}\theta_{source})$

Spot size: $r = \frac{f_X}{F_{FL} \cdot F_{CL}} \left(-\frac{F_2}{F_1} F_{CL} + \frac{f \cdot F_1}{F_2}\right) R_{source} + \frac{f \cdot F_{CL} \cdot F_2}{F_{FL} \cdot F_1} \theta_{source}$

If $L_2 = \frac{F_2}{F_1}(F_1 + F_2)$:

Beam height: $y_2 = -\frac{F_2}{F_1}\left(1 - \frac{L_1}{F_{CL}}\right) y_1 - \frac{F_2}{F_1} F_{CL}\beta_1$ direct rescaling between cases with and without a beam expander

Beam radius: $R^* = -\frac{F_2}{F_1}\left(1 - \frac{L_1}{F_{CL}}\right)R_{source} - \frac{F_2}{F_1}F_{CL}\theta_{source}$ direct rescaling between cases with and without a beam expander

Spot size: $r = \frac{f_X \cdot F_2}{F_{CL} \cdot F_{FL} \cdot F_1}\left(1 - \frac{L_1}{F_{CL}}\right)R_{source} + \frac{f \cdot F_{CL} \cdot F_2}{F_{FL} \cdot F_1}\theta_{source}$

**mfFIFI design and alignment instructions**

The alignment is based on an already existing multi-focal microscope setup, using an excitation MLA or equivalent for generating multi-focal excitation. It is assumed that the front-focal plane of the excitation MLA overlaps with a conjugate image plane.

Due to certain constraints of the system, the easiest alignment procedure is to align all the elements in reverse order.

**1. Aligning the Fourier lens:** Place the Fourier lens so that the distance between the excitation MLA and the Fourier lens is roughly the focal length of the Fourier lens.

**2. Align the flat-fielding MLAs:** Put the flat-fielding MLAs in their respective mounts and mount them into a 30 mm cage system. Fix the position of the second MLA (closer to the Fourier lens) and orient it so that the microlenses are orthogonal to the plane of the light path. Change the orientation of the first MLA to match that of the second MLA. When the two MLAs are misaligned, the resulting field behind the flat-fielding MLAs will cover a smaller surface than if they were aligned. Alignment can also be achieved visually, by looking through the flat-fielding MLAs.

Position the flat-fielding MLAs so that the spacing from the second MLA to the Fourier lens is equal to the focal length of the Fourier lens. Leave enough space to make finer adjustments later.

**3. Align the beam de-expander:** Place the two lenses of the beam de-expander into a telescope configuration, so that they are displaced by the sum of their focal lengths. Place the lens with the longer focal length first, so that the telescope contracts the size of the beam. The two lenses can also be aligned by using a shearing interferometer and assuring that the beam stays collimated after passing through the beam expander. The position of the beam expander with respect to the first flat-fielding MLA is not crucial, but typically placing the two elements closer together is favourable.

**4. Align the rotating diffuser and focusing telescope:** Place the focusing lens and collimating lens into a 30 mm cage system and into a telescope configuration, so that they are displaced by the sum of their focal lengths. This can also be achieved using a shearing interferometer and assuring the beam stays collimated after leaving the telescope. Mark the position of the lenses on the cage system and take out one of the lenses. Slide in the rotating diffuser and put back the missing lens into the telescope configuration. Place the telescope containing the rotating diffuser into the light path so that the collimating lens is facing towards the Fourier lens.

**5. Finely adjust the spot size:** Place a camera in an intermediate image plane or use a dye solution to image the incident excitation onto the camera. Check that the spot size assures diffraction-limited performance on the sample. If not, try moving the rotating diffuser until the spot size improves. If the spot size is still not sufficient, consider using a beam de-expander with a larger contraction factor, or limit the radius of the beam incident on the flat-fielding MLAs by placing an iris.

**6. Finely adjust the excitation pitch:** To finely adjust the pitch of the excitation spots, connect the flat-fielding MLAs with a part of the cage system to fix their distance, and use another part to slide them towards or away from the Fourier lens. Measure the pitch and compare to the desired value dictated by the pitch of the excitation MLA. If the pitch is larger than the desired value, it means that the flat-fielding MLAs are too close to the Fourier lens. If the pitch is too small, then the flat-fielding MLAs are too far away. Adjust the position of the flat-fielding MLAs until the pitch corresponds to the expect value.

### Wave optics simulation platform

Most of the updated simulation platform is based on the previously published software used for widefield flat-field illumination[10].

### Models for PolyE shift in *Chlamydomonas reinhardtii*

We consider the microtubule triplet as a 60 nm structure, with the centers of the A,B and C microtubules separated by $d \cong 20$ nm, forming a $\theta \cong 120°$ angle with respect to the center of the centriole (Supplemental Figure 11a). By projecting the position of the C-microtubule we obtain a radial and a tangential position:

$$d_r = 2d\cos(\theta)$$

$$d_\theta = 2d\sin(\theta)$$

Where $d_r$ is the radial shift and $d_\theta$ the tangential shift.

#### Twist model

In this model, we fix the localization of PolyE to the C-microtubule. We assume a closing of the angle from 120° to 90°, rotating around the inner A-microtubule. This gives:

$$d_r = 2d\,\Delta(\cos(\theta)) = 2d|\cos(120°) - \cos(90°)| \cong 20.0\ \mathrm{nm}$$

$$d_\theta = 2d\,\Delta\sin(\theta) = 2d|\sin(120°) - \sin(90°)| \cong 5.4\ \mathrm{nm}$$

#### Shift and twist model:

In this model we will purely consider the effect of PolyE shifting microtubule localization within the triplet, while maintaining a constant angle of $\theta \cong 120°$. This will result in:

$$d_r = 2d\cos(\theta) = 2d\cos(120°) \cong 20.0\ \mathrm{nm}$$

$$d_\theta = 2d\sin(\theta) = 2d\sin(120°) \cong 34.6\ \mathrm{nm}$$

We can keep the angle constant since the change in angle will not change the initial and final positions of the PolyE localization in this model, since the rotation is happening around the A-microtubule. Considering a broader range of angles at the distal end (90°-120°) would bound the value of the tangential shift between 34.6-40 nm.

### Supplemental Figures

#### Supplemental Figure 1 – mfFIFI development stages.

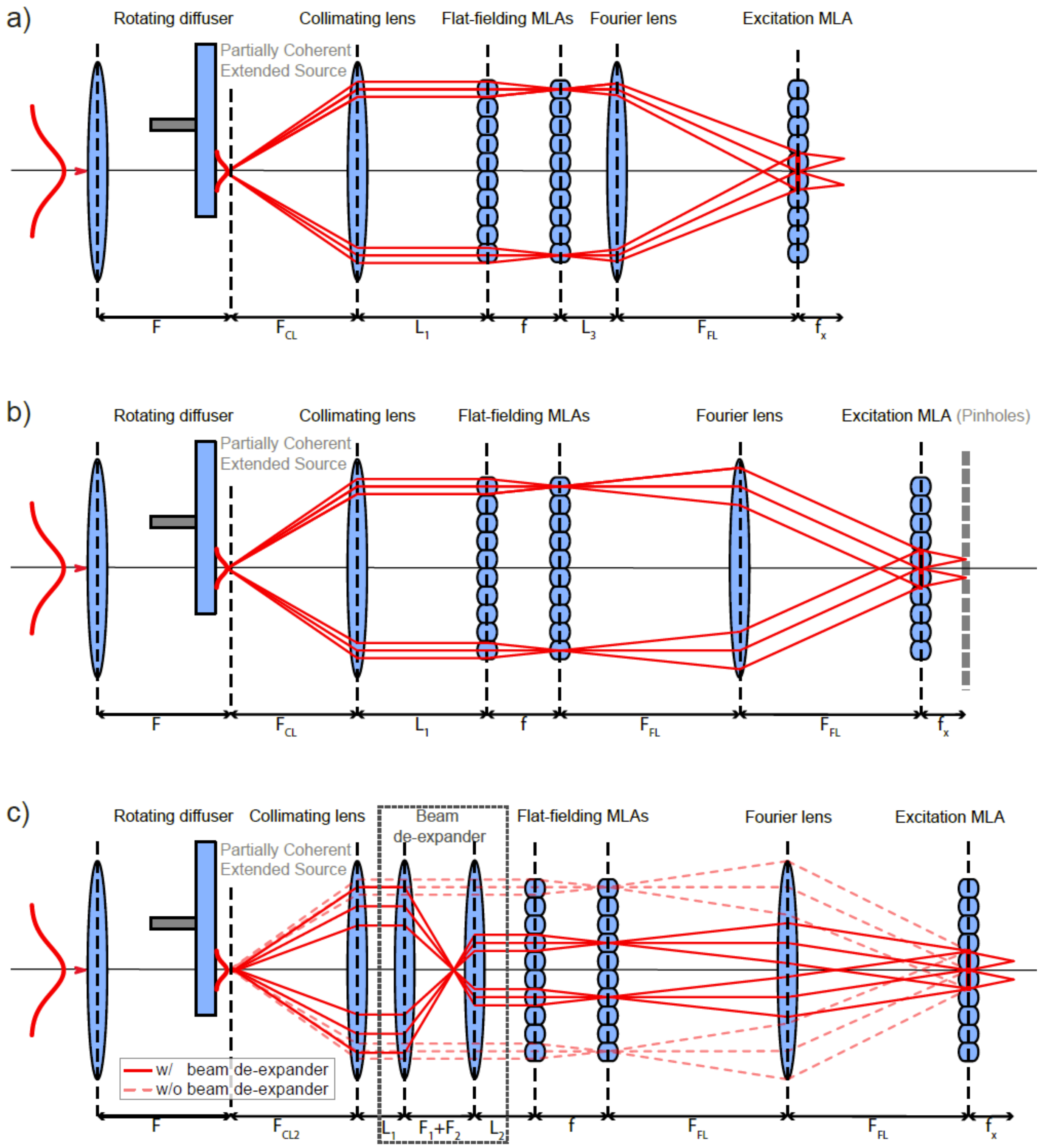


**Supplemental Figure 1 – mfFIFI development stages.** a) A standard Koehler uses a pair of flat-fielding MLAs to split the incoming light into multiple channels, before focusing them in the front-focal plane of the Fourier lens. This effectively averages over spatial variations in the beam intensity, generating a flat-top beam profile. In the case of a coherent light source, such as lasers, a focusing lens and a rotating diffuser are used to scramble the incoming light and generate an extended partially coherent source, which is then collimated by the collimating lens. Implementing a traditional Koehler integrator with a variable length $L_3$ between the second flat-fielding MLA and the Fourier lens, results in non-telecentric illumination of the excitation MLA. This will in turn cause the pitch of the excitation spots produced by the excitation MLA to vary. b) The telecentric Koehler integrator assures that the pitch of the excitation spots generated by the excitation MLA is conserved, by setting the distance between the second flat-fielding MLA and the Fourier lens to the focal length of the Fourier lens: $L_3 = F_{FL}$. Nevertheless, due to the nature of the extended source, the excitation spots will be larger than those produced by direct Gaussian excitation. This will limit the capability of the microscope to achieve diffraction-limited excitation at the sample. A possible solution would be to limit the size of the spots by placing a pinhole array in the front focal plane of the excitation MLA, although at a cost to the transmission efficiency. c) The extended design overcomes this problem by introducing a beam expander between the collimating lens and the first flat-fielding MLA, which allows control over how many microlens channels are used to average over in the focus of the Fourier lens. Careful choice of the beam de-expansion factor enables contraction of the size of the excitation spots, while maintaining good transmission efficiency and homogeneity. Alternatively, placing a hard aperture to limit the radius of the beam incident on the flat-fielding MLAs would have a similar effect, albeit rejecting much of the incident light.

**Supplemental Figure 2 – Ray transfer matrix parameters and coordinates.**

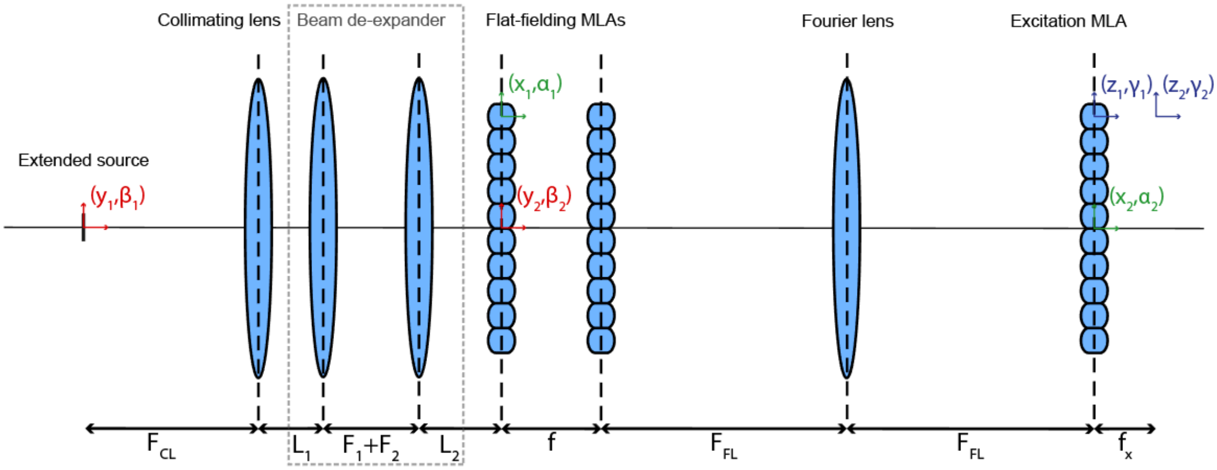


**Supplemental Figure 2 – Ray transfer matrix parameters and coordinates.** Schematic representation of the different coordinate systems used in the ray transfer matrix calculations. The coordinate system $(y_1, \beta_1)$ to $(y_2, \beta_2)$ describes the system from the input beam to the first flat-fielding MLA. The calculation can be performed with and without considering the effect of the beam de-expander. The coordinate system $(x_1, \alpha_1)$ to $(x_2, \alpha_2)$ is used to project the path from each microlens channel to the front-focal plane of the Fourier lens. Finally, $(z_1, \gamma_1)$ to $(z_2, \gamma_2)$ considers the light travelling through each microlens of the excitation MLA, to its front-focal plane.

**Supplemental Figure 3 – Optimization of design parameters using the extended simulation platform.**

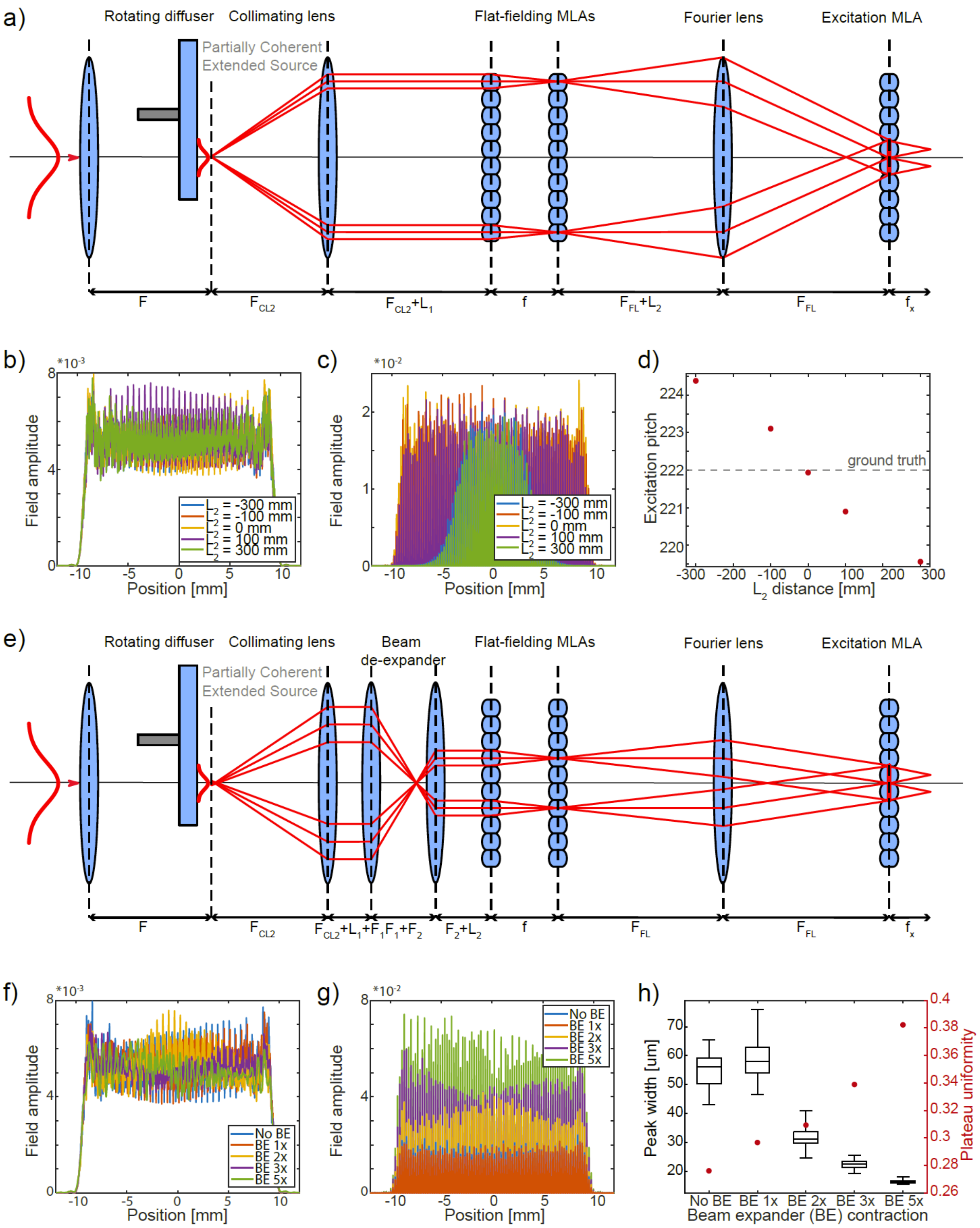


**Supplemental Figure 3 – Optimization of design parameters using the extended simulation platform.** a) Schematic showing the standard Koehler integrator setup, indicating how the different design parameters are defined in the simulation. b) Field amplitude at the front focal plane of the Fourier lens, corresponding to the field incident on the excitation MLA, and c) at the front focal plane of the excitation MLA for different distances $L_2$ of the Fourier lens from the flat-fielding MLAs. The telecentric condition corresponds to $L_2 = 0$. d) Average pitch of the multi-focal excitation measured at the front focal plane of the excitation MLA for different values of $L_2$. Dashed line marks the actual pitch of the excitation MLA. e) Schematic representation showing the mfFIFI configuration including the beam contractor and labeling the simulation parameters. The beam contraction factor is set by the inverse of the magnification of the two lenses of the beam contractor: F2/F1. f) Field amplitude at the front focal plane of the Fourier lens and g) the front focal plane of the excitation MLA. h) Trade-off of the beam contraction factor between the spot size (left axis) and the homogeneity of the excitation spots (right axis), quantified by the plateau uniformity(ref).

**Supplemental Figure 4 – Integrating mfFIFI into an instant structured illumination microscope.**

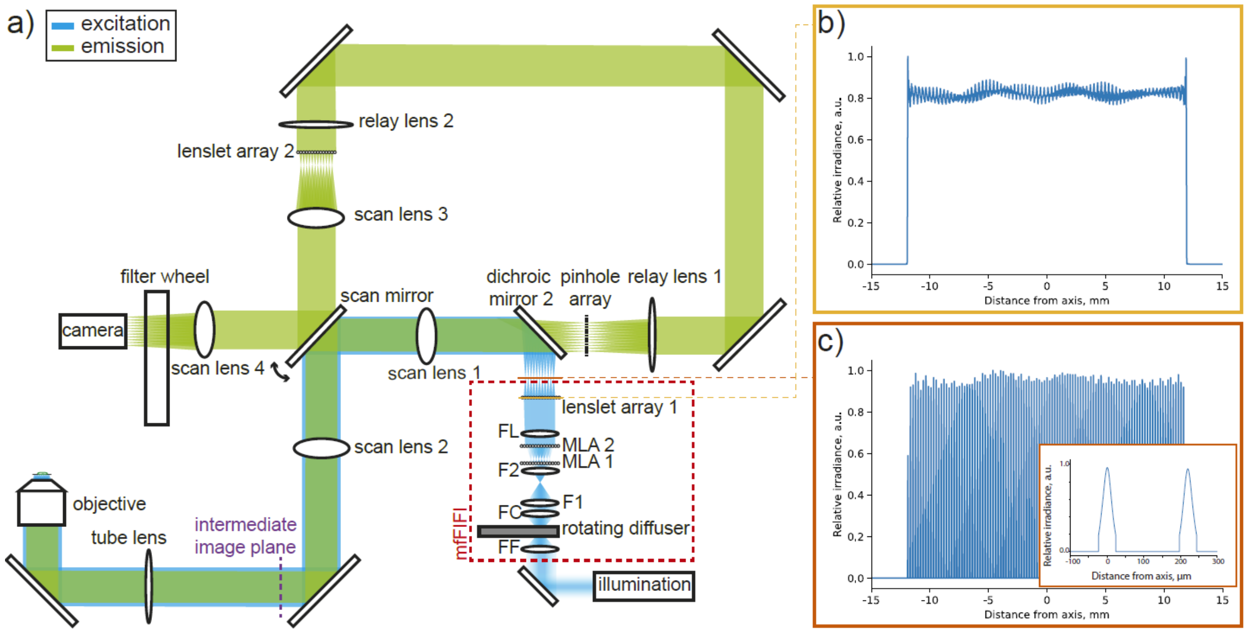


**Supplemental Figure 4 – Integrating mfFIFI into an instant structured illumination microscope.** a) Schematic representation of the iSIM setup, showing where the mfFIFI module is integrated into the excitation path prior to the excitation MLA. b) Simulated flat-field incident on the excitation MLA. c) Simulated intensity profile in the front-focal plane of the excitation MLA, showing an array of excitation points (inset).

**Supplemental Figure 5 – Excitation profiles in iSIM scanning mode.**

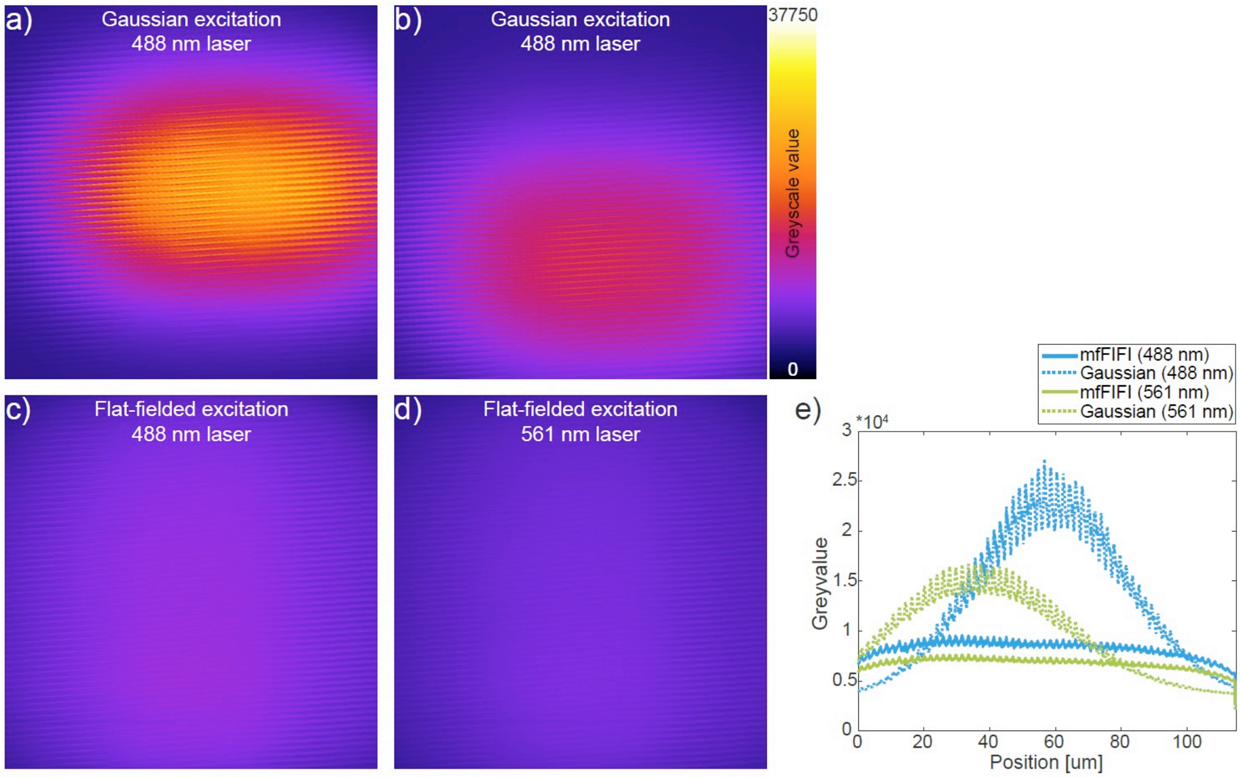


**Supplemental Figure 5 – Excitation profiles in iSIM scanning mode.** a-b) Scanning excitation illumination using Gaussian excitation for the (a) 488 nm and (b) 561 nm lasers using in each case a corresponding fluorescent dye sample. c-d) Scanning excitation illumination using mfFIFI excitation for the (a) 488 nm and (b) 561 nm lasers. e) Intensity profiles along the vertical direction of the different excitation illuminations from (a-d).

**Supplemental Figure 6 – Centriole particles from expanded RPE-1 cells**

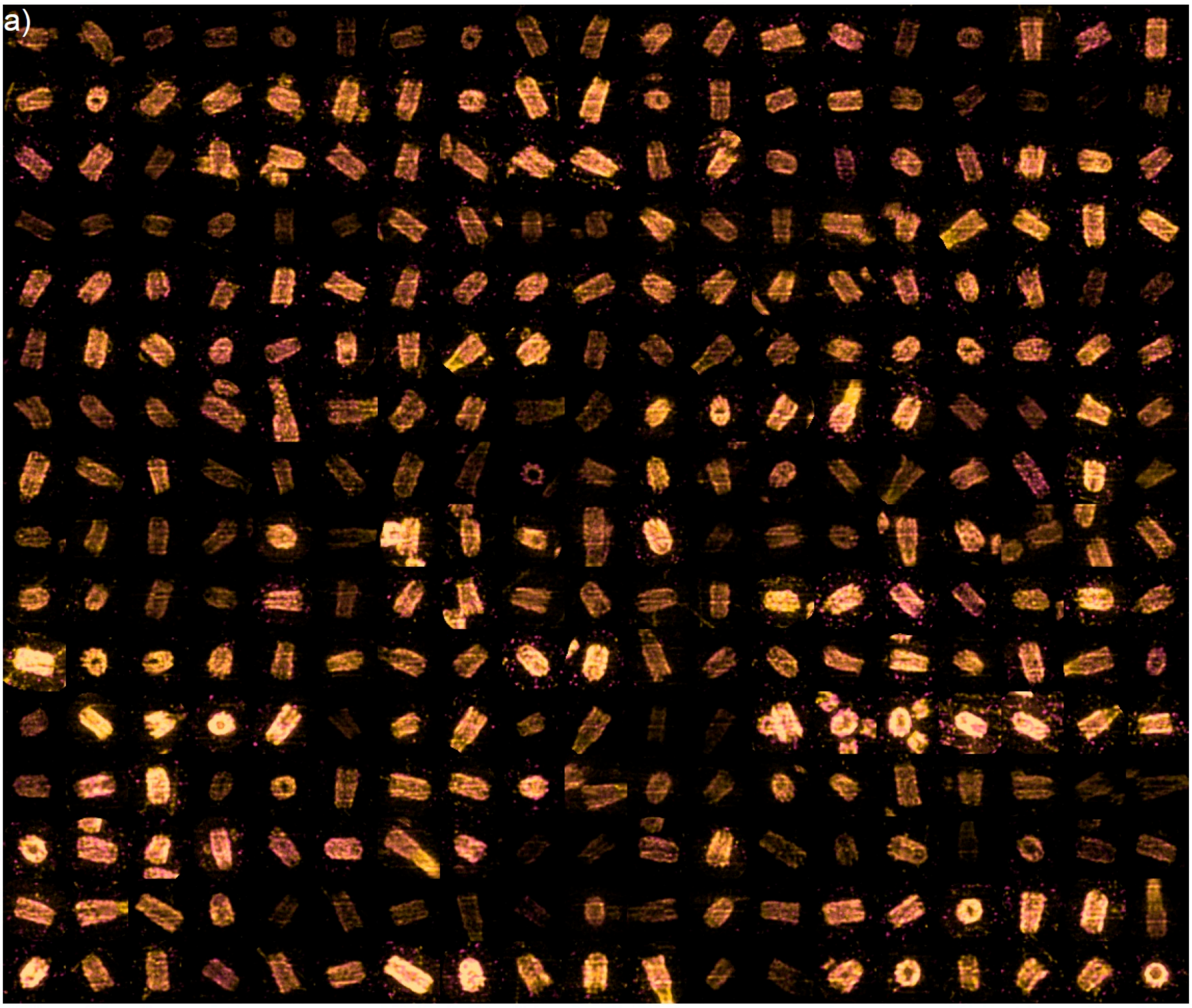


**Supplemental Figure 6 – Centriole particles from expanded RPE-1 cells.** Montage of a random subset of unclassified raw particles collected in situ from expanded synchronized human RPE-1 cells stained for acetylated tubulin (yellow) and PolyE (magenta).

**Supplemental Figure 7 – Particle shape analysis and PTM coverage**

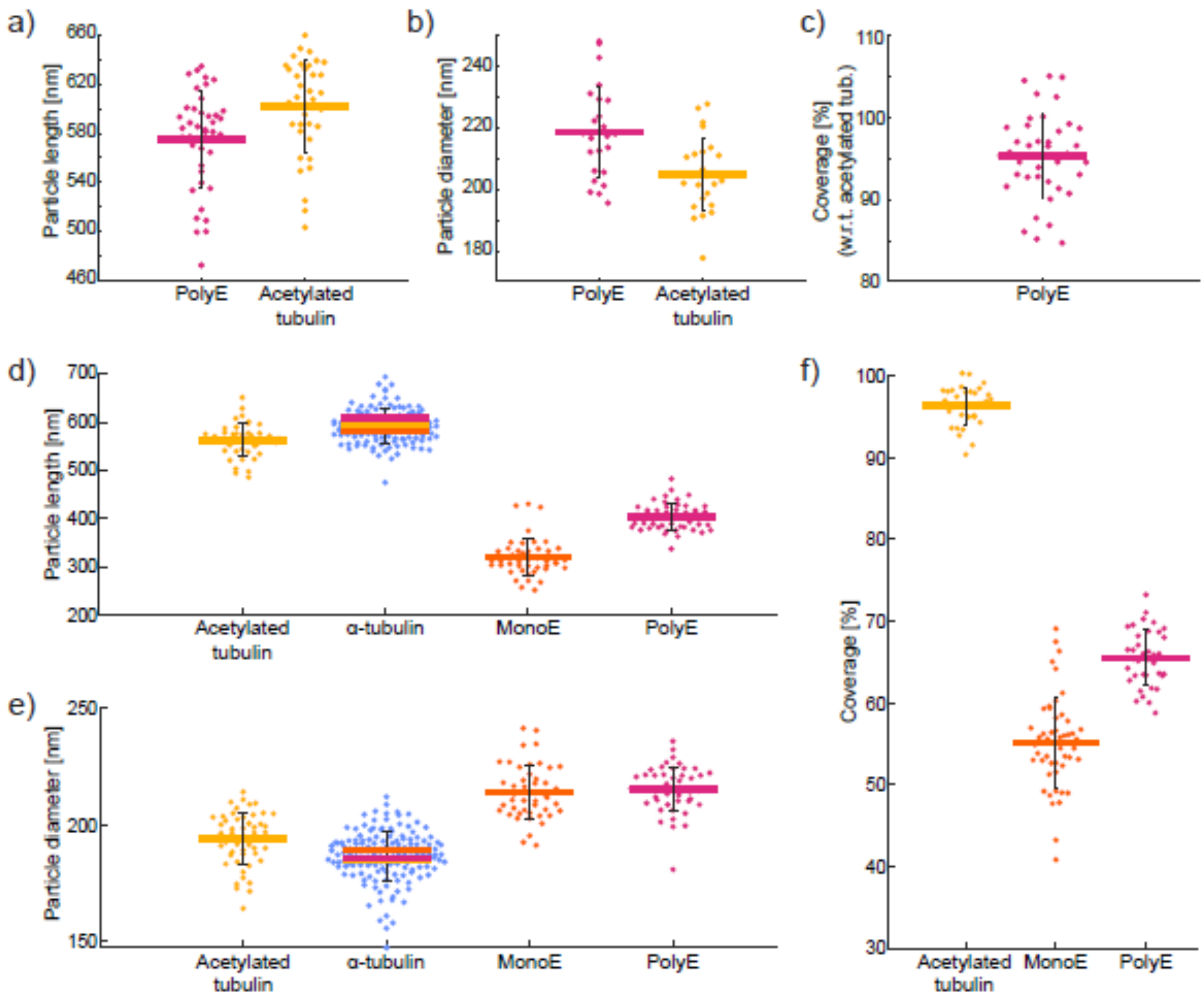


**Supplemental Figure 7 – Particle shape analysis and PTM coverage.** a) Particle lengths measured along side views of expanded human centrioles in the cellular context. b) Particle diameters measured on top views of human centrioles. c) PolyE coverage for human centrioles with respect to the acetylated tubulin signal along the length of the centriole. d) PTM length coverage measured along side views of purified *Chlamydomonas reinhardtii* centrioles. e) Particle diameters measured on top views of purified *Chlamydomonas reinhardtii* centrioles. α-tubulin signal was measured from three different datasets (dual-labeling with acetylated tubulin, MonoE and PolyE), with their individual means marked individually (acetylated tubulin in yellow, MonoE in orange and PolyE in magenta). f) PTM coverage for purified *Chlamydomonas reinhardtii* centrioles measured by dividing the length profiles of different PTMs by their respective α-tubulin signal. All scales reflect pre-expansion size. N = 25 top views and N=41 side views for human centrioles. N = 50 for acetylation dataset, N = 49 for MonoE dataset and N = 50 for PolyE dataset in *Chlamydomonas reinhardtii*. Error bars represent the standard deviation.

**Supplemental Figure 8 – Expanded purified *Chlamydomonas reinhardtii* centrioles**

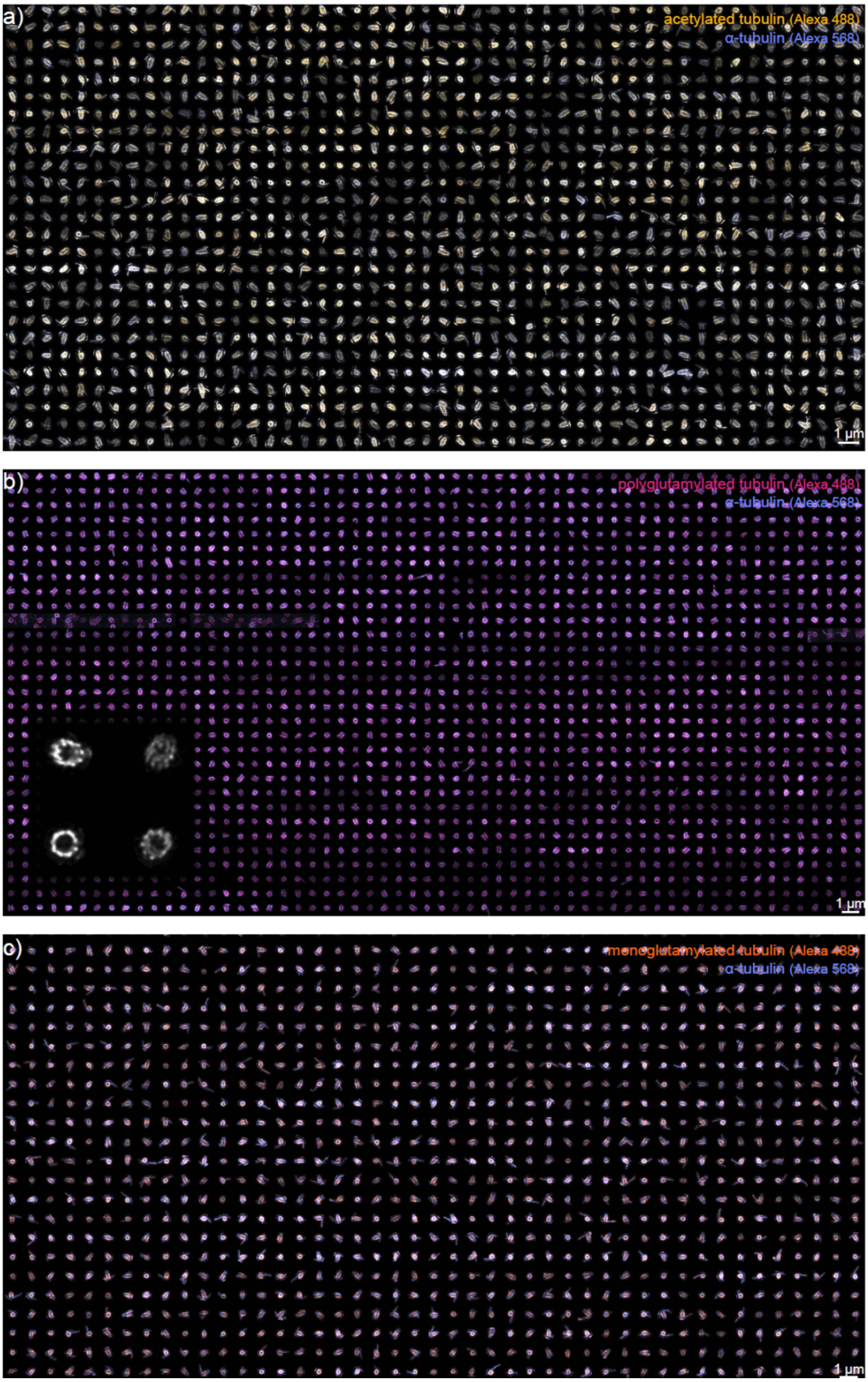


**Supplemental Figure 8 – Particle shape analysis and PTM coverage of purified centrioles.** a-c) Montages of maximum intensity projections of a subset of centriole particles collected with dual-staining for a) acetylated tubulin, b) PolyE and c) MonoE with α-tubulin used as reference in each case. Inset in b) shows examples of PolyE twisting as single color maximum intensity projections.

**Supplemental Figure 9 – Particle classification dendogram**

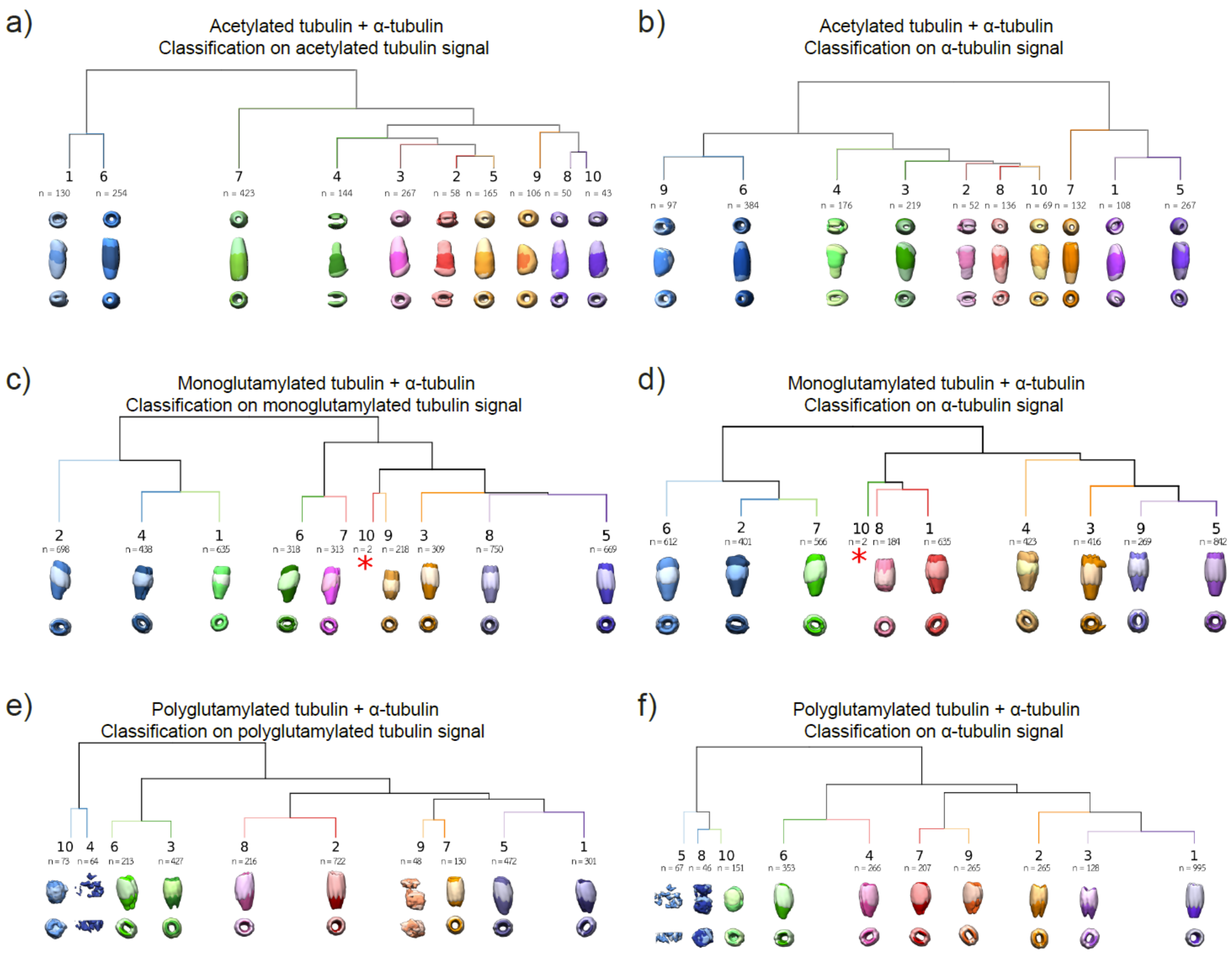


**Supplemental Figure 9 – Particle classification prior to reconstruction.** Dendrograms generated via hierarchical classification in 10 groups. On top, the average volume of all particles is displayed with 2 different orientations. For each group, the average volume is shown with the same 2 orientations, the tubulin signal dark-coloured and the (a-b) acetylated, (c-d) monoglutamylated and (e-f) polyglutamylated signal light-coloured.

**Supplemental Figure 10 – Expanded purified *Chlamydomonas reinhardtii* centrioles**

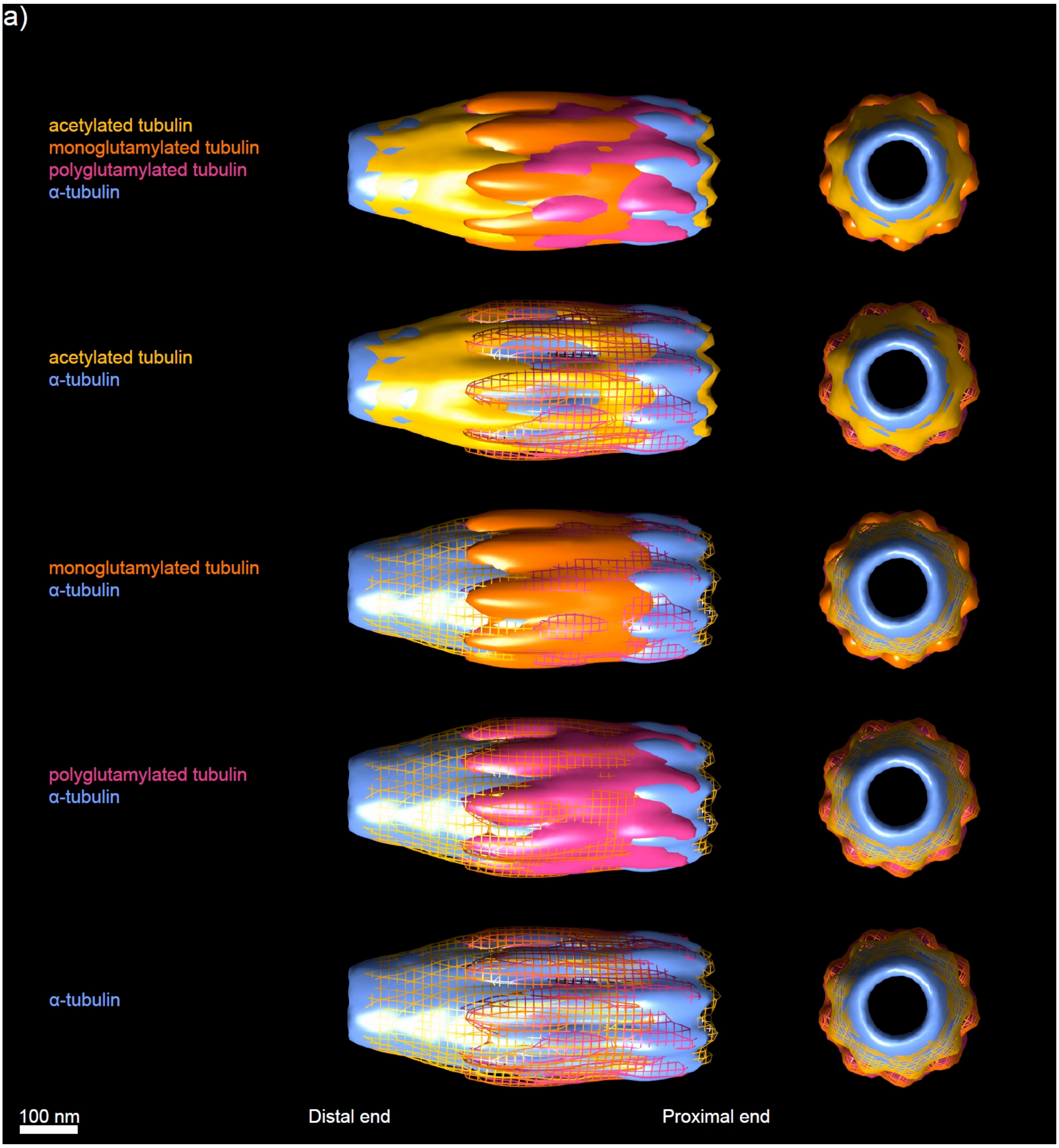


**Supplemental Figure 10 – Multi-color particle averaging and reconstruction of tubulin PTMs in *Chlamydomonas reinhardtii* centrioles.** a) Side and top views of α-tubulin reference with different PTMs. Top views are taken from the distal side toward the proximal side. Scale bar: 100 nm.

**Supplemental Figure 11 – Twist of polyglutamylated tubulin along *Chlamydomonas reinhardtii* centriole**

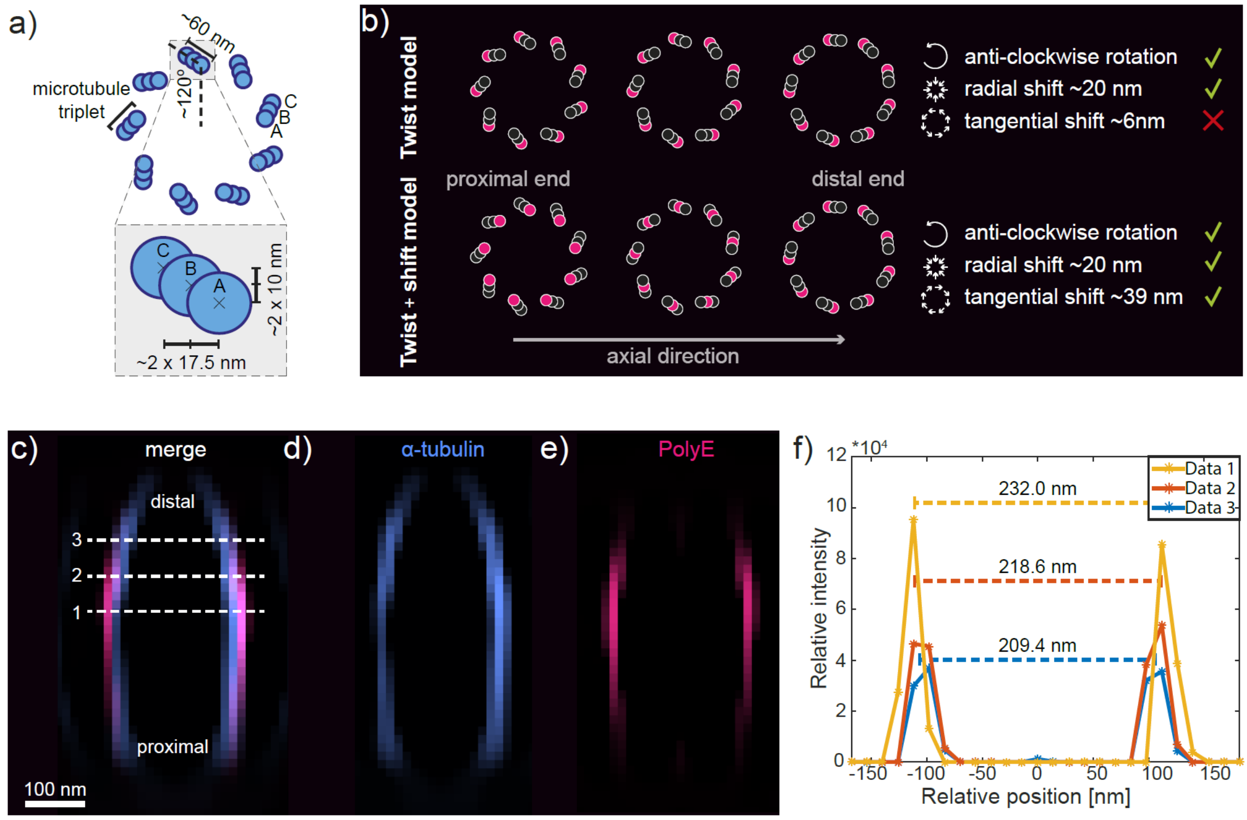


**Supplemental Figure 11 – Twist of polyglutamylated tubulin along *Chlamydomonas reinhardtii* centrioles.** a) Schematic illustration of centriolar microtubule triplets and expected radial and tangential displacements between microtubules in neighbouring triplets (inset) viewed from the proximal end. b) Schematic representation of XY planes between the shift and twist models of PolyE shift along the proximal-distal centriole axis and their predictions (cross sections viewed from the proximal end). c-e) Cross section of the YZ profile showing the barrel diameter with (d) α-tubulin and (e) PolyE signal. f) Intensity profiles measured along the dashed lines from (c) showing the radial displacement of the PolyE signal.

**Supplemental Figure 12 – *Chlamydomonas reinhardtii* reconstruction cross-sections**

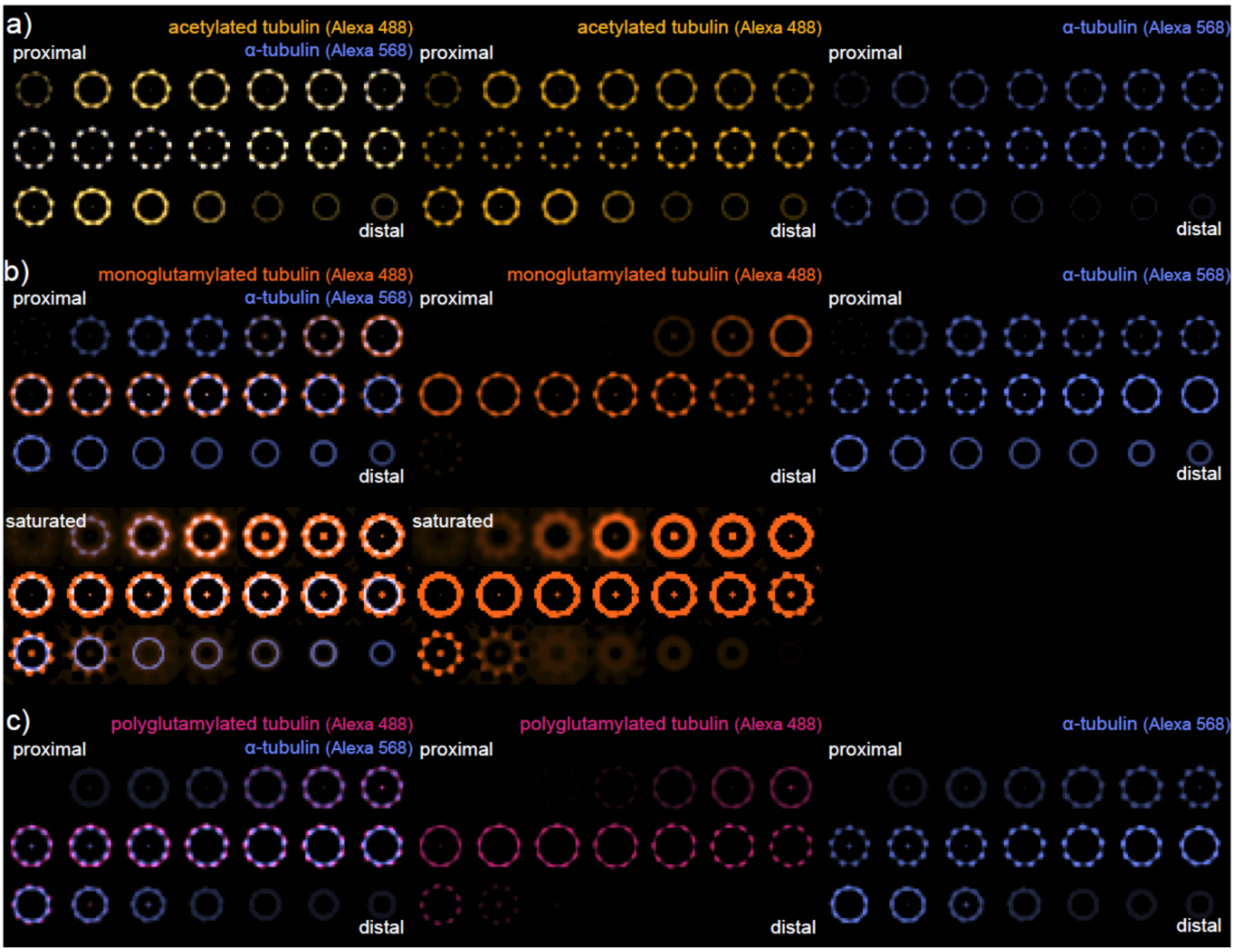


**Supplemental Figure 12 – Chlamydomonas reinhardtii reconstruction cross-sections.** Cross sections starting from the proximal towards the distal end for (a) the acetylated tubulin and α-tubulin, (b) MonoE and α-tubulin and (c) PolyE and α-thubulin datasets. Distance between slices is 28 nm (every second slice from the reconstruction. All images are displayed as viewed from the proximal end. For panels in (b), we also include a second rendering with a saturated intensity scale ("saturated"), to allow coverage at the ends to become more apparent.

### Supplemental Tables

**Supplemental Table 1 – Wave optics simulation parameters**

| Parameters | Simulation 1: effect of $L_2$ (Supplemental Figure 3a-d) | Simulation 2: effect of beam de-expander (Supplemental Figure 3e-h) | Multi-focal excitation test (Supplemental Figure 3) |
|---|---|---|---|
| $\Delta r$, Diffuser offset [mm] | -5000 | -5000 | -5000 |
| $f_{OBJ}$, Objective focal length [mm] | 300000 | 300000 | 550000 |
| $D_{BFP}$, objective BFP diameter [mm] | 50000 | 50000 | 25000 |
| $f_c$, collimating lens focal length [mm] | 60000 | 60000 | 50000 |
| $L_2$, Distance between second MLA and objective BFP [mm] | See figure | 0 | 1000 |
| p, Lenslet periodicity [μm] | 300 | 300 | 1015 |
| N, number of lenslets | 33 | 33 | 11 |
| $f_{MLA}$, lenslet focal length [mm] | 2200 | 2200 | 11000 |
| $L_1$, distance between collimating lens and MLA's [mm] | 5000 | 5000 | 5000 |
| λ, wavelength [μm] | 0.561 | 0.561 | 0.561 |
| $\sigma_f$, diffuser correlation length [μm] | 10 | 10 | 10 |
| $\sigma_r$, diffuser variance | 1.75 | 1.75 | 1.75 |

| | | | |
|---|---|---|---|
| σ, beam standard deviation at waist [μm] | 6 | 6 | 6 |
| nIter, Number of field realizations | 100 | 100 | 1000 |
| Input beam grid size | 50001 | 50001 | 20001 |
| Input beam grid physical size | 50000 | 50000 | 5000 |
| MLA grid size | 2001 | 2001 | 1001 |
| MLA physical grid size | 9900 | 9900 | 11165 |
| BFP grid size | 198099 | 198099 | 660033 |
| $f_1$ beam expander lens 1 focal length | n.a. | 120000 | n.a. |
| $f_2$ beam expander lens 2 focal length | n.a. | 12000, 6000, 4000, 24000 (see figure) | n.a. |

**Supplemental Table 1 – Wave optics simulation parameters.** Parameters used for simulations shown in **Supplemental Figure 3**. Simulation 1 tests the dependence of the spot pitch on the distance $L_2$ between the second flat-fielding MLA and the back focal plane of the Fourier lens (**Supplemental Figure 3a-d**). Simulation 2 tests the dependence of the spot size on the contraction factor of the beam expander introduced between the collimating lens and the flat-fielding MLAs (**Supplemental Figure 3e-h**).

**Supplemental Table 2 – Dataset size and number of particles used for reconstruction**

| Dataset | No. collected particles | No. of particles in reconstruction |
|---|---|---|
| Dataset #1 – *Chlamydomonas reinhardtii* acetylated tubulin (A488) + α-tubulin (A568) | 2129 | 387 |
| Dataset #2 – *Chlamydomonas reinhardtii* GT335 (MonoE) (A568) + α-tubulin (A488) | 6527 | 830 |
| Dataset #3 – *Chlamydomonas reinhardtii* PolyE (A488) + α-tubulin (A568) | 1865 | 910 |
| Dataset #4 – human centrioles PolyE(A488) + acetylated tubulin (A568) | 392 | - |

**Supplemental Table 2 – Dataset size and number of particles used for reconstruction.** Numbers of segmented particles obtained from the raw data of purified centrioles and used for particle averaging and reconstruction. The segmentation had a ~50% efficiency and improving the segmentation pipeline could possibly extract more particles from the same raw datasets.
